# Cotranslational assembly confers specificity for in vivo target heterodimerization of paralogous H2B-like TAF12 proteins in the human fungal pathogen *Candida albicans*

**DOI:** 10.1101/2025.06.23.660276

**Authors:** Vidhi Bhardwaj, Selene Swanson, Laurence Florens, Michael P. Washburn, Jerry L. Workman, Krishnamurthy Natarajan

**Affiliations:** Laboratory of Eukaryotic Gene Regulation, School of Life Sciences, Jawaharlal Nehru University, New Delhi, India; Stowers Institute for Medical Research, Kansas City, MO, USA; Department of Cancer Biology, University of Kansas Medical Center, Kansas City, KS, USA

## Abstract

The fidelity of assembly of multiprotein complexes is essential for the formation of stable and functional protein complexes that are critical for cell growth and survival. In this context, TBP-associated factor (TAF) subunits maintain tight specificity for their integration into TFIID and SAGA complexes. In this work, using affinity purification-coupled mass spectrometry of epitope-tagged TFIID subunits TBP and TAF11, and the SAGA subunit TAF12L we identified components of the *C. albicans* TFIID and SAGA complexes. Whereas TAF12 is a subunit of TFIID, the paralogous TAF12L is a subunit of the SAGA complex, and we further identified each of the TFIID and SAGA complex subunits with high confidence. We found that the steady-state levels of the H2B-H2A-like histone fold domain containing pairs, TAF12-TAF4 and TAF12L-Ada1 proteins, are mutually dependent on the stable expression of each other. Using RNA coimmunoprecipitation from polysome-containing extracts, we found that nascent TAF4 and Ada1 proteins interact with TAF12 and TAF12L, respectively, by a cotranslational mechanism in an ordered, sequential mode of assembly. Thus, our results indicate that heterodimerization of the TAF12 paralogs with cognate partners occur by sequential cotranslational assembly thereby ensuring both selectivity and stability of the H2A-H2B heterodimers in fungal pathogen *C. albicans*.

## Introduction

Transcriptional complexes are all large, heteromeric, and module-containing megadalton complexes. It is a prerequisite for the transcriptional factors to be assembled correctly to regulate gene expression. The complexity of large multisubunit assembly requires an authentic identification of the partners to avoid non-specific interactions that could turn lethal for the organism. Hurdles are expected when considering the problem of correct polypeptide recognition and folding while forming oligomeric structures so that they function competently.

Although a protein’s folding pathway might vary based on several conditions, a common set of rules governs the process of assembly and folding for various proteins. One such idea is cotranslational assembly, which posits that a protein starts folding and assembly begins even as the ribosome synthesizes the polypeptide chain. Both eukaryotic and prokaryotic cells have devised the mechanism of cotranslational assembly, where protein subunits of a complex assemble while at least one of the interacting partners is at the nascent stage of being translated at the ribosome. The process of cotranslation provides faithful recognition of subunits, which is hard to expect if the multiprotein assembly were to occur by mere random collision.

The way one or more nascent polypeptides interact with their cognate partners while they are synthesized at the polysomes defines which kind of co-translational assembly is occurring. When two polypeptides interact at their nascent stage while being synthesized, it is called a co-co type of cotranslation. This can be either through cis interaction, i.e., polypeptides translated from one mRNA exiting the polysome, or trans, i.e., polypeptides translated from different mRNAs. The co-co assembly is prevalent in prokaryotic systems for the formation of homo/heteromers. The cis-assembly is exclusive for homomers and prominent in bacteria. The operon system in bacteria enables the formation of heterodimers translated from polycistronic mRNA. The assembly of LuxA and LuxB subunits of bacterial luciferase provides a fitting example (1). In eukaryotes, heterodimer formation is only possible through trans-assembly. How the nascent polypeptides find each other is still poorly understood. However, smRNA-fish (single-molecule fluorescence in situ hybridization) provides visualization of mRNAs present in proximity in vivo. A study show that the cotranslationally assembled Rpt1 and Rpt2 mRNA co-localized together in Not1-assemblysomes, whereby the co-localized RNA not only assembled but also prevented the aggregation process. These assemblysomes are believed to provide necessary factors for ribosome pausing, folding, and translation elongation(2). Another example of co-co assembly is TAF6-TAF9 heteromers-assembly(3). The other mode is the sequential mode of cotranslational assembly-called the Co-post type. Briefly, an already-made polypeptide that has exited the ribosome tunnel interacts with its partner at the nascent stage of its synthesis when the interacting interface is exposed. Examples are TAF8-TAF10, Set1C, Nucleoporins, among many (3–5). Dimers also assemble hierarchically around one subunit, acting as a scaffold and driving the complex assembly. For example, TAF1, the largest subunit of the TFIID complex, is enriched in almost all the RNA-immunoprecipitations performed for other TAFs (6).

The human fungal pathogen *C. albicans* is categorized among the top fungal pathogens in the critical fungal pathogen list and has two TAF12 paralogs, TAF12 and TAF12L. In this study, we utilized MudPIT analysis of TAF12 and TAF12L containing complexes and showed that TAF12 paralogs are exclusive for SAGA or TFIID. We aimed to dissect whether the association of both these Histone-fold-containing TAF12 paralogs occurs through the cotranslational assembly pathway as a mechanism to impart stability and selectivity. We report that both TAF12L-ADA1 and TAF12-TAF4 heterodimerization occur through the cotranslational mechanism sequentially and are specific for nascent polypeptide binding.

## Results

### MudPIT analysis revealed that TAF12 and TAF12L paralogs are unique subunits of TFIID and SAGA in *C. albicans*

In a previous study using coimmunoprecipitation assays from cell extracts, we reported that the *C. albicans* TAF12 paralogs, TAF12 and TAF12L, interact with TFIID and SAGA complexes, respectively (7). To further understand the TAF12 and TAF12L interactions with TFIID and SAGA, we employed Multidimensional Protein Identification Technology (MudPIT) mass spectrometry analysis of immunopurified protein complexes. Towards this end, to affinity purify TFIID, we used *C. albicans* strains expressing TBP-TAP and TAF11-TAP strains. The TAP-purified complexes were eluted, and mass spectrometry analysis was carried out. To affinity purify SAGA complex, we used a 3xFLAG-tagged TAF12L strain or SN87 as an untagged control and carried out affinity purifications using anti-FLAG M2 affinity agarose, protein complexes eluted, and mass spectrometry analysis was carried out. We analysed the MudPIT results for proteins showing significant enrichment, based on dNSAF scores and peptide coverage, for the known TFIID and SAGA subunits as per the genome annotations in the Candida Genome Database (8).

We identified homologs of each of the subunits of the *S. cerevisiae* SAGA complex (9, 10) in the TAF12L-FLAG purified sample, including the shared TAFs viz., TAF5, -6, -9, -10, and -12 (Fig. 1*B*). However, none of the other TFIID subunits were identified in our preparation (Fig. 1; Table S3). In the TAF11-TAP and TBP-TAP purified samples, homologs of each of the yeast TFIID subunit including TAF12, but no TAF12L was identified (Fig. 1*B*; Table S3). These results showed that TAF12L and TAF12 are subunits of the *C. albicans* SAGA and TFIID complexes, respectively. Our study also revealed several proteins that interact with TBP, TAF11, and TAF12L, in addition to the bonafide TFIID and SAGA subunits, raising the possibility of identifying novel interacting proteins (Fig. 1*A*, Table S3). Thus, unlike in *S. cerevisiae*, where TAF12 is shared in both TFIID and SAGA complexes, TFIID and SAGA in *C. albicans* contain two paralogous TAF12 proteins.

**Figure 1.**
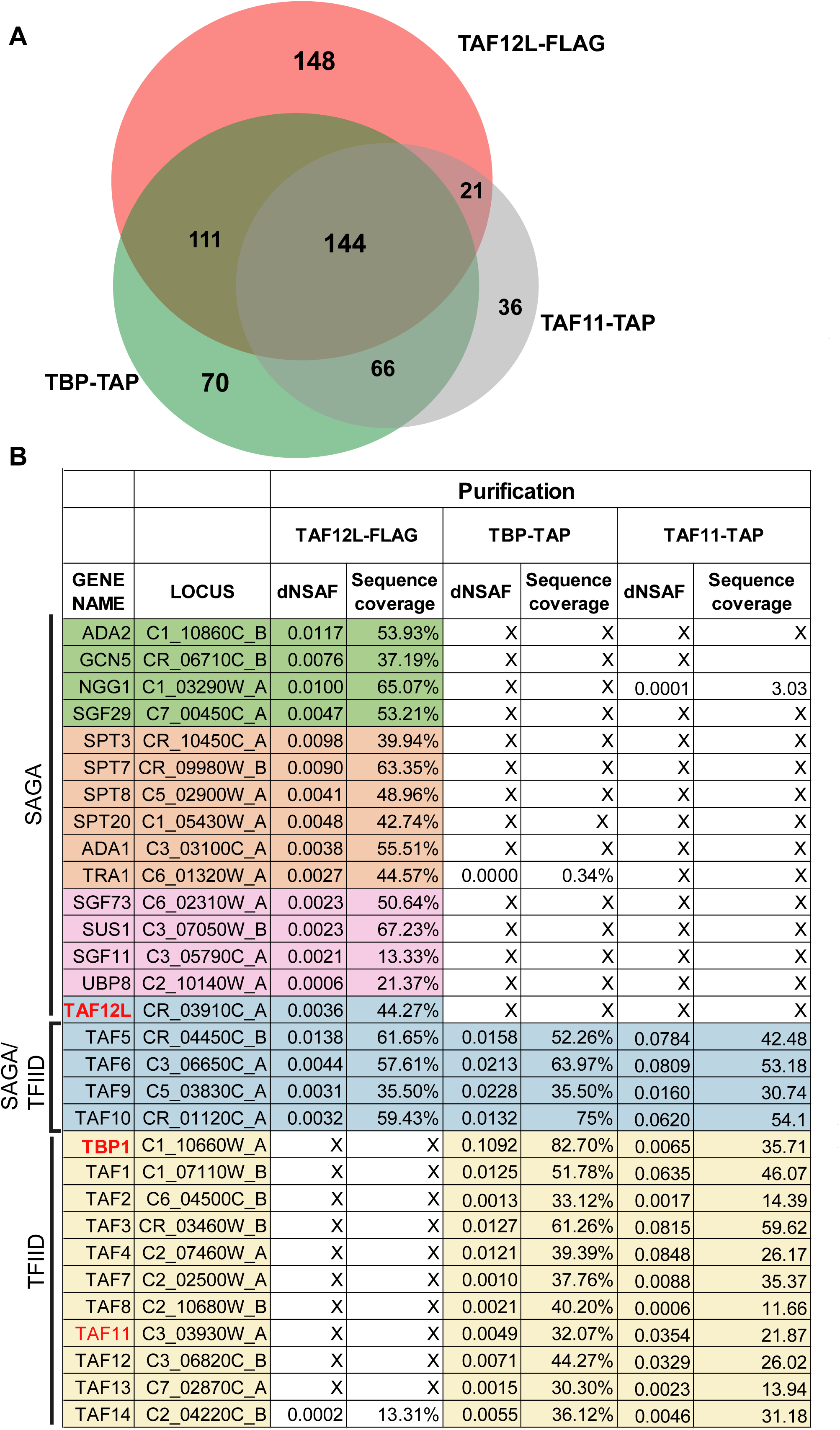
MudPIT identification of *C. albicans* TFIID and SAGA. *A,* Venn diagram showing the TFIID and SAGA subunits enriched upon immunopurification-coupled mass spectrometry of TAF12L-FLAG, TBP-TAP, and TAF11-TAP purifications. *B,* List of TFIID and SAGA subunit proteins identified MudPIT data. The distributed Normalized Spectral Abundance Factor (dNSAF) and percentage sequence coverage for each protein are shown. The homologs of protein subunit names were obtained from Candida Genome Database annotation; the asterisks indicate that the protein names are assigned in this study as per homology to the *S. cerevisiae* proteins.

### Interdependence of TAF12L-ADA1 and TAF12-TAF4 for protein stability

Our genetic analysis showed that *C. albicans TAF12* is essential for cell growth, and *TAF12L*, although not essential for growth, but was required for maintaining wild-type yeast cell morphology in *C. albicans* (11). The histone H2B-like histone fold domain (HFD) of TAF12 forms a heterodimer with H2A-like HFD-bearing Ada1 or TAF4 proteins in human and yeast (12, 13). It has been reported that the interacting subunits become unstable when the dimerization partner is removed in cells (14), or when expressed alone in bacterial cells (15), as the unbound subunit(s) are thought to become unstable as they are prone to misfolding. To test the impact of depletion on the heterodimerizing partner, we constructed conditional depletion mutants by replacing the promoters of *TAF12*, *TAF4*, *TAF12L* or *ADA1* genes with the maltose-regulatable promoter *P_MAL2_* as described previously (11). The promoter-replaced strains ISC11 (*P_MAL2_-TAF12L*), VBC6 (*P_MAL2_-ADA1*), ISC12 (*P_MAL2_-TAF12*), VBC4 (*P_MAL2_-TAF4*) and SN95 (WT) were cultured overnight in YP medium containing maltose, diluted into fresh YP medium containing glucose, cells harvested at 0h, 2h, 4h, and 6h, and protein levels were examined by Western blot analysis. The data showed that under conditions of depletion, TAF12L depletion led to reduction in Ada1 protein levels by 2h (Fig. 2*A*, lane 6) and similarly Ada1 depletion diminished TAF12L protein levels by 4 h (Fig. 2*B*, lane 7). In the parental control strain, however, both TAF12L and Ada1 protein levels remain unchanged at all time points ((Fig. 2*A-B*, lanes 1-4). Next, we examined the level of TAF4 and TAF12 proteins upon TAF12 (ISC12) or TAF4 (VBC4) depletion in the respective strains. We observed that the TAF4 protein level was undetectable upon TAF12 depletion (Fig. 2*C*, lane 7), and reciprocally, TAF12 level was significantly diminished upon TAF4 depletion (Fig. 2*D*, lane 8). In parental control strain, both the level of TAF12, TAF4 remain unchanged (Fig. 2*C*-*D*, lanes 1-4). We also tested the growth phenotypes of *TAF4* (VBC4) and *ADA1* (VBC6) depletion mutants and found that *TAF4* is essential for cell growth, while *ADA1* was not required for growth in YPD (Fig. S1). To examine if the effect of depletion observed at protein levels was due to transcriptional downregulation of their respective mRNAs, we examined the level of *TAF12*, *TAF4, TAF12L*, and *ADA1* mRNAs post 6h of depletion. We found that the mRNA levels encoding the heterodimerizing partners do not change significantly upon depletion (Fig. 2*E-F*). Together, these results showed that for protein stability heterodimerizing partners require each other. These data suggests that the heterodimeric proteins provide chaperoning function to each other to maintain protein stability indicating a likely cotranslational mechanism for subunit recognition thus providing high specificity and stability to the heterodimers as observed in other cellular systems (16–18).

**Figure 2.**
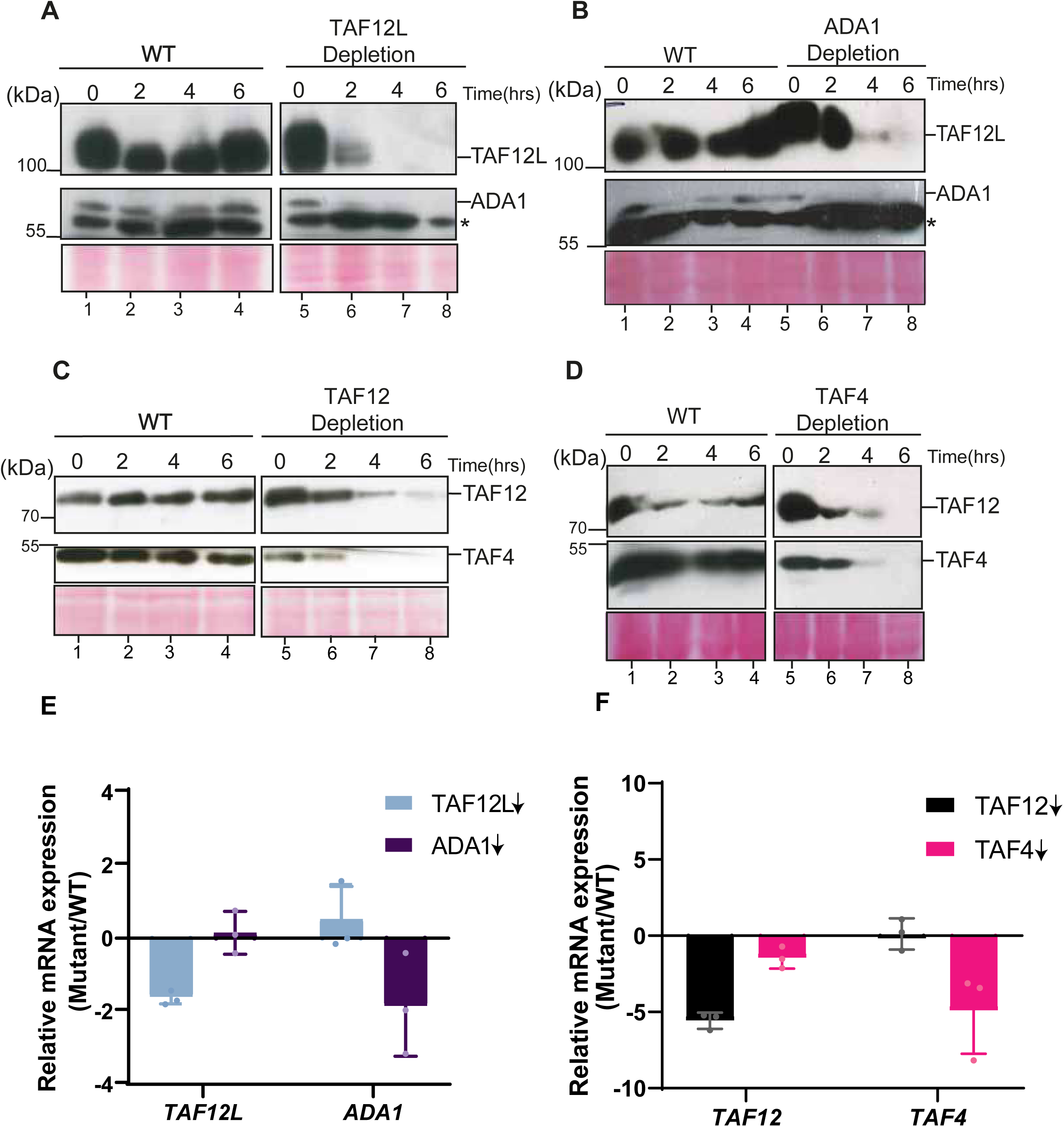
Protein stability of the TAF12-TAF4 and TAF12L-Ada1 heterodimers is dependent on each partner. *A-D,* Western blot analysis of cell extracts from *A,* ISC11 upon TAF12L (*P_MAL2_*-*TAF12L*) depletion. *B*, VBC6 upon Ada1(*P_MAL2_*-*ADA1*) depletion, *C,* ISC12 upon TAF12L (*P_MAL2_*-*TAF12*) depletion, *D,* VBC4 upon TAF4 (*P_MAL2_*-*TAF4*) depletion, or (*A-D*, Left) from SN95 (WT) at indicated time points from 0h to 6h after shift from YP + maltose (*P_MAL2_* ON) to YP + glucose (*P_MAL2_* shut-off). The lower band in the blot probed with anti-Ada1 is a cross-reacting band marked with an asterisk (*). The Western blot membranes were probed with the following antibodies: polyclonal anti-TAF12L (1:1000), anti-Ada1(1:1000), anti-TAF12(1:1000), anti-TAF4 (1:1000), or anti-TAP (1:3000) antibodies. (*E-F*) mRNA levels of indicated transcripts upon *TAF12L*, *ADA1*, *TAF12* or *TAF4* depletion (marked with down arrow). Total RNA was extracted from WT, ISC11, ISC12, VBC4, and VBC6 strains after culturing for 6 h in YPD at 30°C. qPCR analysis was carried out, and relative mRNA expression level (mutant/WT) was plotted, normalized to *SCR1* RNA as endogenous control.

### TAF12 paralogs associate with their cognate heterodimeric partners in a cotranslational manner

To test whether TAF12 paralogs utilize a cotranslational mechanism for heterodimer recognition, we performed RNA-immunoprecipitation followed by qPCR as previously reported (3–5, 19). The mRNA enrichment in protein-protein interaction analysis is indicative of a nascent polypeptide association via translating ribosomes. Therefore, we prepared polysome-containing cell extracts with translation elongation inhibitor-cycloheximide (CHX) to stall translation and stabilize the RNA-protein transient via ribosome(20). Our initial attempts to employ CHX for translational arrest was unsuccessful (data not shown) at standard CHX concentrations employed in other systems, including the budding yeast (Fig. S2) (21). Another study reported polysome profile from *C. albicans* treated with a high concentration of 1mg/ml CHX for polysome analysis (22). Therefore, we first tested *C. albicans* cell growth at different CHX concentrations from 0.2mg/ml to 2 mg/ml, and found that *C. albicans* growth was not significantly impaired till ∼1 mg/ml CHX (Fig. 3*A*). Next, we prepared polysome extracts from cells treated with CHX (1 mg/ml) or untreated cells, and carried out 10%-50% sucrose density gradient centrifugation, and fractions were monitored at A260nm. The polysome profile showed an accumulation of polysomes in CHX-treated cell extracts compared to ribosomal run-off found in the untreated cells (Fig. 3*B*). These results showed that CHX can be effectively employed for translational arrest to make polysome-containing cell extracts.

**Figure 3.**
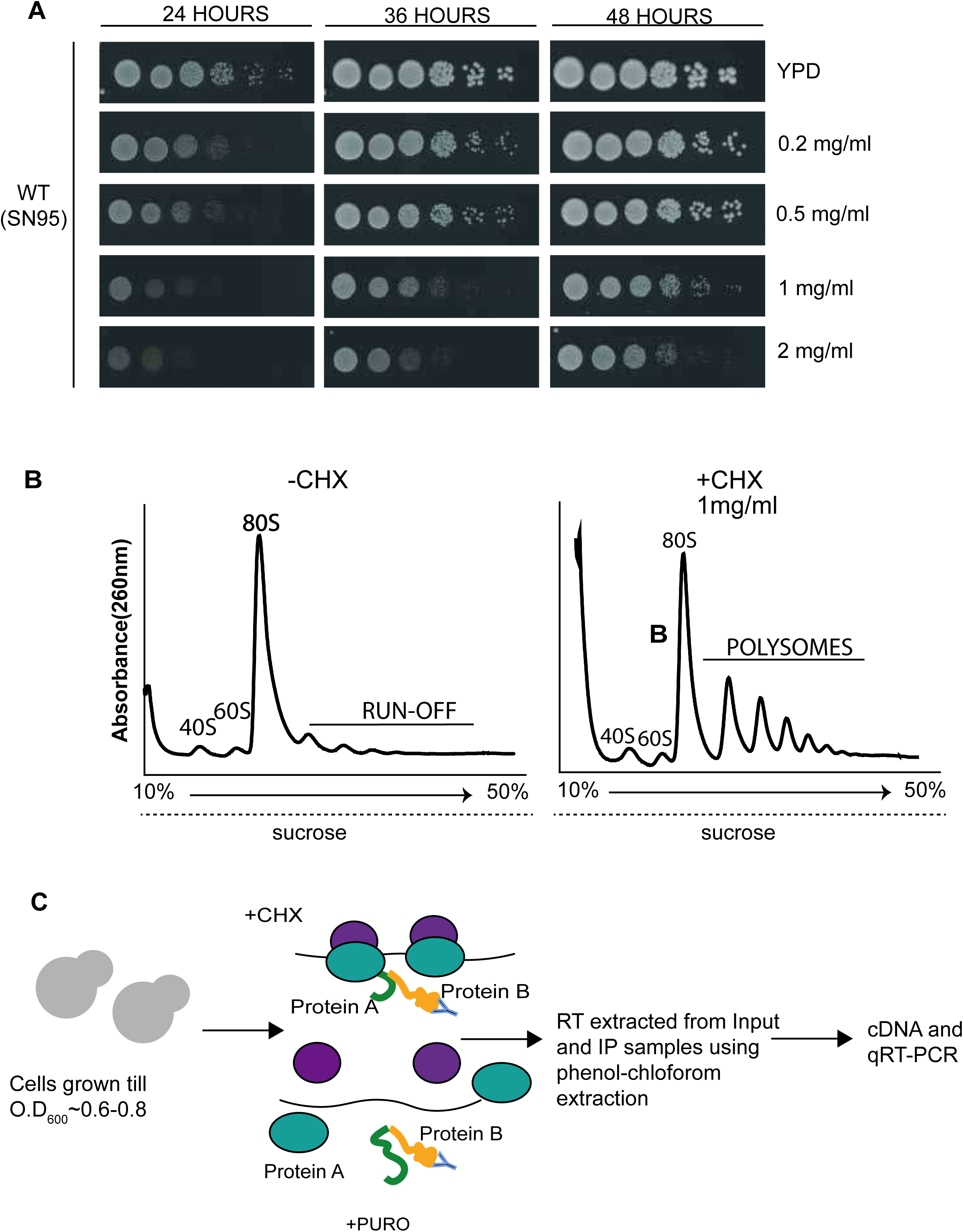
Schema of RNA immunoprecipitation assay using polysome extracts in *C. albicans*. *A*, Optimization of cycloheximide (CHX) concentration on *C. albicans.* WT strain (SN95) was grown in YPD for 16-18h, serially diluted and spotted on YPD, or YPD containing 0.2mg/ml, 0.5mg/ml, 1mg/ml, or 2mg/ml cycloheximide, and incubated at 30°C and photographed at indicated times. *B,* Polysome profiles of cell extracts from CHX-treated or untreated *C. albicans* cultures fractionated through a 10% to 50% sucrose gradient. *C*, Schematic diagram of the RNA-immunoprecipitation (RIP) protocol.

We next set up RIP assays using polysome-containing cell extracts as schematically shown in Fig. 3*C*. *C. albicans* cells were grown in YPD, and either treated with 1mg/ml CHX before harvest, or untreated cells as a control. Polysomal cell extracts were prepared from both CHX-treated and untreated cells to ensure complete ribosomal run-off, the untreated control cell extracts were treated with puromycin. TAF12L-FLAG was immunoprecipitated with anti-FLAG M2 beads from both CHX- and puromycin-treated cell extracts and the coprecipitated RNA were extracted, simultaneously RNA was also extracted from input sample, and used for RT-qPCR analysis. The TAF12L RIP strongly enriched the *ADA1* mRNA in CHX-treated conditions compared to puromycin-treatment, whereas *TAF4* or other mRNAs, i.e., *TAF12L’s* own mRNA, control *SCR1* RNA or *RPS8A* mRNA did not show any significant enrichment difference between the CHX and puromycin conditions (Fig. 4*A*). These results indicated that TAF12L binds to nascent ribosome-associated Ada1 thereby enriching *ADA1* mRNA (Fig. 4*A*). Conversely, RIP with C-terminally tagged Ada1-FLAG as bait showed no enrichment of *TAF12L*, *ADA1*, *TAF12*, control *SCR1* RNA, or the *RPS8A* mRNA in CHX or puromycin-treated conditions (Fig. 4*B*). Taken together, the RIP analysis of both TAF12L and Ada1 proteins indicated that the fully translated TAF12L having exited the ribosome binds to nascent Ada1 polypeptide and, as a result, pulls down ribosome-associated *ADA1* mRNA.

**Figure 4.**
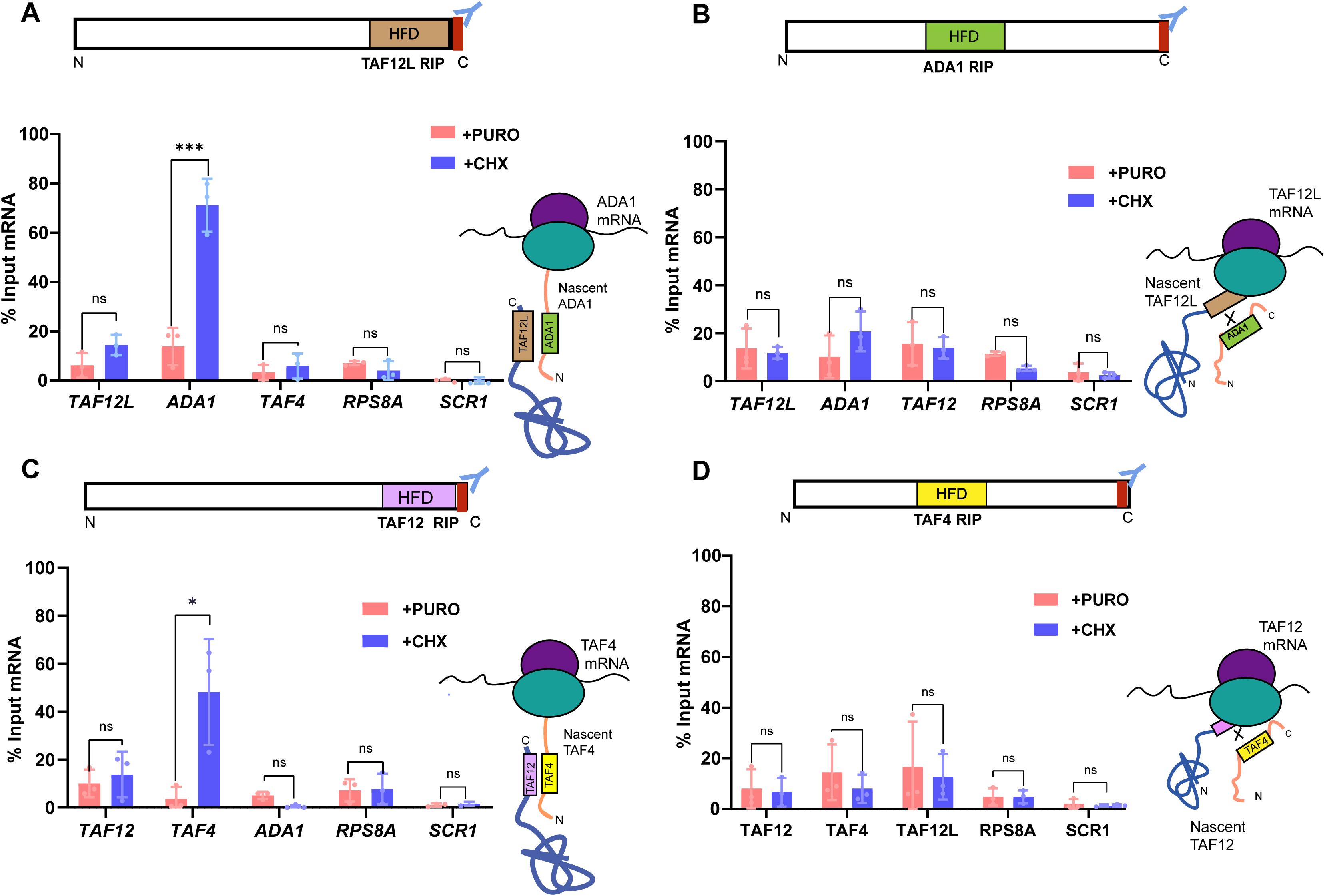
Cotranslational association of Taf12 paralogs and their heterodimers: *A,* RIP performed with Taf12L-FLAG-tagged protein as bait from polysome extract treated with either cycloheximide (CHX) or Puromycin (PURO). *B*, RIP performed with the Ada1-FLAG-tagged protein. *C*, RIP with TAF12-FLAG, and *D*, RIP with TAF4-FLAG. The mRNA enrichment was calculated for each mRNA after normalization corresponding input values, separately for puromycin- and cycloheximide-treated samples, and plotted as percent input mRNA. The values were determined from three biological replicates each, and the error bars represent SD. The significant difference between CHX and PURO-treated samples for each mRNA was determined by applying a student’s t-test. *p*-value ≤ 0.05 (∗), ≤0.01 (∗∗), ≤0.001 (∗∗∗), and >0.05 (ns).

Next, to examine whether TAF12 interacts with TAF4 also in a cotranslational manner, we carried out RNA-immunoprecipitation from TAF12-FLAG containing polysomal extracts as described above. The RIP qPCR analysis showed strong enrichment of *TAF4* mRNA from CHX-treated but not from puromycin-treated cell extracts (Fig. 4*C*). Other mRNAs that we tested alongside, i.e., *ADA1*, *TAF12*, and the control *SCR1* RNA and *RPS8A* mRNA, were not enriched from either of the treated cell extracts (Fig. 4*C*). In contrast, TAF4-FLAG RIP did not enrich *TAF12* mRNA and other RNAs tested (Fig. 4*D*). These results indicate that TAF12 interacts cotranslationally with its partner TAF4 and not vice versa. Taken together, our results support the sequential cotranslational assembly model for TAF12-TAF4 and TAF12L-Ada1 heterodimerization based on the position of the interacting domains along the polypeptide chain. The TAF12 and TAF12L histone fold domains, located near the C-terminal region, facilitates association with nascent TAF4 and Ada1, respectively, bearing HFD towards their N-terminal region, thus enabling an early-stage interaction during ongoing translation. Additionally, an early stage recognition of heterodimerizing partner confers stability to the interacting pair (17).

## Discussion

A concerted, correct assembly of protein complexes is crucial for proper functioning to regulate key processes critical for cell survival. The binding of protein partners in their fully translated state, i.e., post-translationally, exhibits several disadvantages in a crowded cellular environment, such as non-specific associations. As the *C. albicans* TAF12 and TAF12L proteins share ∼77% sequence similarity and ∼57% sequence identity between their histone-fold domains (7), such high degree of sequence similarity could potentially lead to promiscuous cross-interactions, such as TAF12L-associating with TAF4 or TAF12-associating with Ada1. However, our findings depict that the TAF12 paralogs maintain tight specificity driven by the cotranslational assembly mode for partner protein recognition. To prevent non-specific associations in crowded cellular environments, cotranslation of interacting proteins has emerged as a robust mechanism that not only prevents non-canonical association but also prevents misfolding of proteins by stabilizing the partner protein, a condition that otherwise could lead to detrimental consequences.

Several studies have identified numerous cotranslational assembled proteins from transcription factors to nucleoporins and cytoskeletal proteins, implicating that the extensiveness of cotranslational assembly is not limited to certain classes of proteins (2–5, 23). Cotranslational assembly mediate hierarchical assembly for efficient complex formations for large multimeric proteins such as the nucleoporin assembly in *S. cerevisiae*, reducing the time at each step which cannot be attained through simple diffusion of interacting proteins (5). Despite advantages and extensiveness, cotranslational assembly is not an obligatory mechanism, same protein may assemble with distinct partners through different assembly mechanisms based on the outcome of the protein complex for function and interactions, for instance, TAF10 heterodimerization with TAF8 utilized co-translational assembly in TFIID, but same TAF10 did not assemble with SAGA-specific SUPT7 through a cotranslational mode (3, 24). Another such example are the proteins involved in nucleoporins, Seh1 interacts cotranslationally with Nup85 but not with Sea-complex(5).

In this study, we focused on the two TAF12 paralogs in the human fungal pathogen *C. albicans* and have shown that the assembly of both the paralogs occurs through a cotranslational pathway where the nascent TAF4 assembled with TAF12 and this association enriched TAF4 mRNA in a translational-dependent manner, similarly nascent Ada1 assembled with TAF12L and thus enriched ADA1 mRNA. Both the nascent interactions were sensitive to puromycin treatment. Both TAF12 and TAF12L are associated in a sequential mode of assembly because the interacting histone-fold domain is located towards the C-terminus of the TAF12 paralogs. The antipodal locations of the interacting histone fold domain on these interacting TAFs give them a spatiotemporal advantage for the association to happen during ongoing translation. In the sequence of assembly for these pairwise associations, in SAGA, TAF12L was likely synthesised before its partner Ada1, and in TFIID, TAF12 is likely synthesised before TAF4. Since, TAF12-TAF4 and TAF12-Ada1 heterodimers were cotranslationally assembled, we show that a failure of heterodimerization upon depletion, led to degradation of unbound protein. However, the mRNA expression of heterodimers upon depletion did not alter, hinting at a quality control mechanism for protein degradation in the absence of one binding partner, possibly due to exposure of certain residues on the orphaned subunits, which target the protein for degradation (14, 25). A persistently unbound and unfolded protein can cause cellular toxicity. In the context of the pathogen, such an observation could serve the purpose of inhibiting fungal pathogens as excluding one partner, targets the degradation of key transcriptional and functional proteins important for cell survival (26). Understanding the extent of cotranslationally interacting proteins and their regulation becomes increasingly important, especially in identifying key essential transcriptional factors that govern fungal pathogenesis, as targeting these proteins for degradation could become a novel approach to inhibiting fungal invasiveness. Our study will set a basis for identifying other cotranslationally assembled proteins/complexes in fungal pathogens.

## Materials and methods

### Strains and growth conditions

*C. albicans* strains SN87 and SN95 were used as parental strains. All strains were cultured in yeast extract-peptone (YP) rich medium with either glucose or maltose as carbon source, as indicated. The *C. albicans* genome sequence data and annotations were obtained from the Candida Genome Database (CGD). Details of strain construction are described in the supplemental text. The list of strains used are provided in Table S1, and the oligonucleotides used in Table S2.

### Immunoblotting

Indicates strains were cultured in YPM at 30°C for 16-18 hours with shaking at 220 rpm, and diluted the culture into fresh YPD and collected cells at 0, 2, 4, and 6 hours by centrifugation. Whole-cell extracts were made by bead beating using chilled glass beads for 8 cycles in lysis buffer as described (27). Quantified proteins using Bradford assay (Bio-Rad) using BSA as a standard. 100µg was loaded into each well of 8% SDS-PAGE. The blots were probed with 1:1000 dilution of anti-TAF12, TAF12L, TAF4, and Ada1 polyclonal antibodies. All blots were visualized using the ECL Plus chemiluminescent system.

### RNA extraction and RT-PCR

*C. albicans* strains were grown as described under Immunoblotting. About 10 O.D_600_ were harvested by rapid filtration, and RNA was extracted using the hot-phenol method (28). One microgram of RNA was subjected to DNase I treatment, and one-third was converted to cDNA using Superscript III cDNA synthesis kit using random hexamer primers. RT-qPCR was performed with 4μl of 1:50 diluted cDNA as template and gene-specific primers (Table S2) in duplicate for each sample. The relative mRNA levels were calculated by ΔCq (WT)-ΔCq (mutant), where ΔCq=Cq(gene)-Cq (*SCR1*), and ΔΔCq values were plotted using GraphPad Prism 8.0 as relative mRNA levels (Mutant/WT).

### Polysome extract preparation and polysome analysis

*C. albicans* strain SN95 was cultured overnight till saturation in YPD, diluted to fresh YPD, and cells were grown till OD_600_ ∼0.6-0.8, and 1mg/ml CHX was added to the culture for 5 min, and cells from 100 ml were harvested, typically total ∼70 OD_600_. The cell pellets were washed with lysis buffer (20 mM HEPES KOH pH 7.5, 150 mM KCl, 10 mM MgCl_2,_ 0.1% (vol/vol) NP-40, and 1x Roche Complete protease inhibitor) containing with or without 1mg/ml CHX, resuspended in ∼200μl of the same buffer with or without CHX and lysed by glass bead lysis for 10 min each using Vortex Genie instrument. Cell extracts containing 5 A260 units were loaded onto a 12ml 10-50% (wt/vol) sucrose gradient and centrifuged in a SW41Ti rotor at 35000 rpm for 160min at 4°C in Beckman Coulter ultracentrifuge. At the end of the centrifugation, samples were fractionated using an automated gradient maker and fractionator (BioComp Gradient Master and Fractionator, USA), and A260nm of the fractions were recorded and plotted.

### RNA-Immunoprecipitation

The polysome extracts were prepared as described above except that the cell pellets were washed with 10 ml lysis buffer containing either 1mg/ml CHX (+CHX culture) or 1mg/ml puromycin (+Puro culture), and cells again resuspended in ∼600μl of the lysis buffers with CHX or Puro, and 40U RNase inhibitor (Thermo Fischer) was added to each sample just prior to the extract preparation. Cells were lysed in presence of glass beads and vortexed for 10 minutes, and ∼4mg equivalent total proteins from whole cell extracts were used for immunoprecipitation using 30μl anti-FLAG M2 affinity gel (Sigma/Merck) for 4h at 4°C. The beads were then washed four times with the washing buffer (lysis buffer with 350mM KCl), and beads resuspended in AE buffer (50 mM NaOAc, 10 mM EDTA) with 20μl 10% SDS per sample, extracted with an equal volume of pre-warmed phenol, followed by chloroform extraction, and the aqueous phase was collected, and ethanol precipitated in presence of NaOAc and 1μl glycogen. The pellet was washed, dried, and dissolved in 10μl nuclease-free water (Ambion). Ten percent of the input cell extract from each sample was used for RNA preparation as input RNA control as above. Five-microliters of 1:10 diluted input RNA and 5 μl of immunoprecipitated RNA were treated with DNase I and used for cDNA preparation, and qPCR was carried out as described above. Quantitative enrichment analysis was carried out using the formula 100 × 2^[(Cq(Input)−3,322)−Cq(IP)]^ and expressed as % Input RNA as described previously (24).

### Affinity Purification and LC-MS/MS analysis

*C. albicans* TAF12L-FLAG (SKC3), TBP-TAP (ISC33), TAF11-TAP (ISC49), and control (SN87) strains were precultured for 14h in YPD and then diluted into six litres of YPD and grown till mid-log phase. The cells were collected by centrifugation, washed, and resuspended in either 20ml H350 buffer (25mM HEPES-KOH, pH=7.5, 350mM KCl, 2mM MgCl2, 1mM EDTA, 10% glycerol, 0.02% NP40) for FLAG samples, or 12ml TAP extraction buffer (40 mM HEPES-KOH, pH 7.5, 10% Glycerol, 350mM NaCl, 0.1% Tween-20) with protease inhibitors (1mM PMSF, 1μg/ml pepstatin A, 2μg/ml leupeptin, and 100μl Sigma Yeast Protease Inhibitor (P8250) per 50ml cell suspension). The cells were lysed in a BeadBeater at 4°C, and heparin (0.5mg for FLAG extracts, or 1mg for TAP extracts) and 125 units benzonase were added, centrifuged at 45000 rpm for 1.5h, and the supernatants were collected. For FLAG purification, 400μl pre-washed anti-FLAG M2 affinity agarose gel (A2220, Sigma) was added to the SKC3 and SN87 lysates, and incubated with rotation at 4°C for 3-5h. The beads were washed, and bound proteins were eluted by the addition of 250μg/ml 3xFLAG peptide (F4799, Sigma), incubated for 30 min at 4°C, and elution repeated 3-4 times and fractions collected.

For purification of TAP-tagged complexes, two-step purification was employed as outlined briefly. The lysates from TAP-tagged (ISC33 and ISC49) and untagged control (SN87) strains were incubated with 400μl IgG-Sepharose that was pre-washed with TAP extraction buffer and incubated for 2h on a rotator. The beads were collected by gentle centrifugation at 600-1200 rpm, and the supernatant was separated. The bead fractions were washed five times with TAP extraction buffer at 4°C, and bound proteins were eluted by TEV cleavage as follows. The beads were incubated with 1ml TEV cleavage buffer containing 10μl AcTEV (Invitrogen), incubated with rotation at 4°C for 16h, and the eluates were collected. Next, the eluate was added to 250μl pre-washed Calmodulin-Sepharose beads in calmodulin binding buffer (CBB; 10mM Tris, pH 8.0, 1mM MgOAc, 1mM Imidazole, 2mM CaCl_2_, 0.1% NP-40, 10% Glycerol, 0.3M NaCl and protease inhibitors) containing 1M CaCl_2_ and incubated at 4°C for 3h on a rotator, and the beads washed five times with CBB (0.15M NaCl). The bound proteins were eluted with 200μl calmodulin elution buffer (10mM Tris, pH 8.0, 150mM NaCl, 1mM MgOAc, 1mM imidazole, 2mM EGTA, 0.1% NP-40, 10% Glycerol, protease inhibitors), serially seven times, and the fractions were collected. The eluted samples were precipitated using 20% (w/v) TCA, washed with cold acetone, air-dried and submitted for LC-MS/MS mass spectrometry.

### Multidimensional Protein Identification Technology (MudPIT)

TCA-precipitated protein pellets were resuspended in 30 µl of 100mM Tris-HCl, pH 8.5, 8 M urea, reduced with 5mM TCEP (Tris(2-Carboxylethyl)-Phosphine Hydrochloride, Pierce), and alkylated with 10 mM IAM (Iodoacetamide, Sigma). As described (29), a two-step digestion procedure was used. Endoproteinase Lys-C (Roche) was added to 0.5 µg for at least 6 hours at 37°C, then the sample was diluted to 2M urea with 100mM Tris-HCl, pH 8.5. Calcium chloride was added to 2mM and the digestion with trypsin (0.5 µg) was let to proceed overnight at 37°C while shaking. The reaction was quenched by adding formic acid to 5% and the peptide mixture was loaded onto a 100µm fused silica (FS) microcapillary column packed with 8 cm of reverse phase material (Aqua, Phenomenex) and connected to a 250µm FS column packed with 3 cm of 5-μm Strong Cation Exchange material (Partisphere SCX, Whatman), followed by 2 cm of 5-μm C_18_ reverse phase(30).

The loaded microcapillary column was placed in-line with a Quaternary Agilent 1100 series HPLC pump. Overflow tubing was used to decrease the flow rate from 0.1 ml/min to about 200–300 nl/min. Fully automated 10-step chromatography runs were carried out (31). Three different elution buffers were used: 5% acetonitrile, 0.1% formic acid (Buffer A); 80% acetonitrile, 0.1% formic acid (Buffer B); and 0.5M ammonium acetate, 5% acetonitrile, 0.1% formic acid (Buffer C). Peptides were sequentially eluted from the SCX resin to the reverse phase resin by increasing salt steps, followed by an organic gradient. The last two chromatography steps consisted in a high salt wash with 100% Buffer C followed by the acetonitrile gradient. The application of a 2.5 kV distal voltage electrosprayed the eluting peptides directly into a LTQ linear ion trap mass spectrometer equipped with a nano-LC electrospray ionization source (Thermo, San Jose, CA). Full MS spectra were recorded on the peptides over a 400 to 1,600 *m*/*z* range, followed by five tandem mass (MS/MS) events sequentially generated in a data-dependent manner on the most intense ions selected from the full MS spectrum (at 35% collision energy). Mass spectrometer scan functions and HPLC solvent gradients were controlled by the Xcalibur data system (Thermo, San Jose, CA). SEQUEST(32) was used to match MS/MS spectra to peptides in a consisting of 9269 non-redundant proteins derived from the all orf translation of Candida albicans SC5314 Assembly 22(http://www.candidagenome.org/download/sequence/C_albicans_SC5314/Assembly22/current/; as described (33), 354 usual contaminants (such as human keratins, IgGs, and proteolytic enzymes), and, to estimate false discovery rates, 9623 randomized amino acid sequences derived from each non-redundant protein entry. The validity of peptide/spectrum matches was assessed using the SEQUEST-defined parameters, cross-correlation score (XCorr) and normalized difference in cross-correlation scores (DeltCn). Spectra/peptide matches were only retained if they had a DeltCn of at least 0.08 and, minimum XCorr of 1.8 for singly-, 2.0 for doubly-, and 3.0 for triply-charged spectra. In addition, the peptides had to be fully-tryptic and at least 7 amino acids long. Combining all runs, proteins had to be detected by at least 2 such peptides, or 1 peptide with 2 independent spectra. Under these criteria the final FDRs at the protein and peptide levels were less than 1%. DTASelect (30)was used to select and sort peptide/spectrum matches passing this criteria set. Peptide hits from multiple runs were compared using CONTRAST(30). To estimate relative protein levels, spectral counts were normalized as distributed Normalized Spectral Abundance Factors (dNSAFs) as described (34).

## Data Availability

Supplemental material containing figures and methods are provided; all strains and plasmids are available upon request. Original data underlying this manuscript generated at the Stowers Institute can be accessed after publication from the Stowers Original Data Repository at http://www.stowers.org/research/publications/libpb-2562.

## Author Contributions

VB and KN conceptualized the study, performed the experiments, analysed data, and drafted the manuscript. All proteomics purifications were carried by KN at the Stowers Institute, and MudPIT analysis carried out by SS, and MW at the Stowers Institute Proteomics facility. JLW and KN acquired funding, supervised the project, and finalized the manuscript.

## Acknowledgments

We thank the members of the Natarajan lab for discussions; KN thanks Swaminathan Venkatesh and Shanshan Li for advice and discussions, and Laszlo Tora for discussions and advice; Leos Valasek, and Kaustuv Datta for advice on polysome analysis. We thank Anil Thakur and Nidhi Gupta for assistance and the use of the instruments for polysome analysis at the BRIC-Regional Centre for Biotechnology, Faridabad. This study was supported by a grant from DST-SERB (EMR/ 2017/000161; CRG/2022/005145) and through departmental funding from DBT-BUILDER (BT/INF/22/SP45382/2022) and DST FIST-II (SR/FST/LSII-046/2016(C) program grants.

## Supplementary information

**Table S1.** List of strains and plasmids.

| Strain | RELEVANT GENOTYPE | SOURCE |
| --- | --- | --- |
| SN95 | *arg4Δ/arg4Δ his1Δ/his1Δ URA3/ura3Δ::imm<sup>434</sup> IRO1/iro1Δ::imm<sup>434</sup>* | (Noble and Johnson, 2005) |
| SN87 | *ura3Δ-iro1Δ::imm434/URA3-IRO1, his1Δ/his1Δ, leu2Δ/leu2Δ* | (Noble and Johnson, 2005) |
| ISC11 | SN95 *HAH1-P<sub>MAL2</sub>-TAF12L/HIS1-P<sub>MAL2</sub>-TAF12L* | (Sinha et al., 2017) |
| ISC12 | SN95 *HAH1-P<sub>MAL2</sub>-TAF12/HIS1-P<sub>MAL2</sub>-TAF12* | (Sinha et al., 2017) |
| SKC3 | SN87 *TAF12L::His6-Gly2-FLAG3-FRT/TAF12 L::His6-Gly2-FLAG3-SAT1-FLP* | (Sinha et al., 2017) |
| SKC6 | SN87 *TAF12:: His6-Gly2-FLAG3-FRT/TAF12 ::His6-Gly2-FLAG3-SAT1-FLP* | (Sinha et al., 2017) |
| SKC8 | SN87 *ADA1::His6-Gly2-FLAG3-FRT/ADA1* | (Sinha et al., 2017) |
| SKC12 | SN87 *TAF4::His6-Gly2-FLAG3-FRT/TAF4::His6-Gly2-FLAG3-SAT1-FLP* | (Sinha et al., 2017) |
| VBC3 | *SN95 P<sub>MAL2</sub>TAF4/TAF4* | This work |
| VBC4 | *SN95 P<sub>MAL2</sub>TAF4/P<sub>MAL2</sub>TAF4* | This work |
| VBC5 | *SN95 P<sub>MAL2</sub>ADA1/PADA1* | This work |
| VBC6 | *SN95 P<sub>MAL2</sub>ADA1/P<sub>MAL2</sub>ADA1* | This work |

**Table S2.**
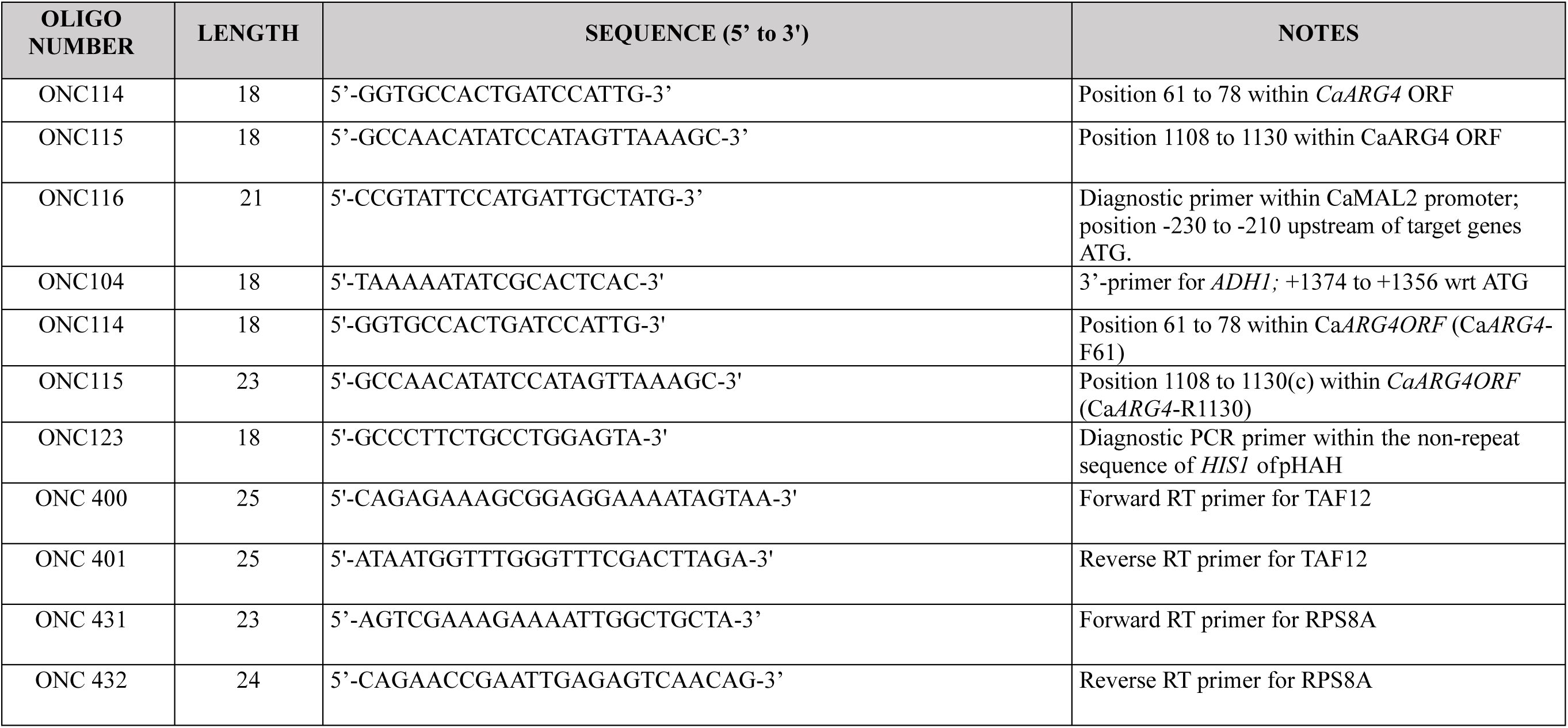
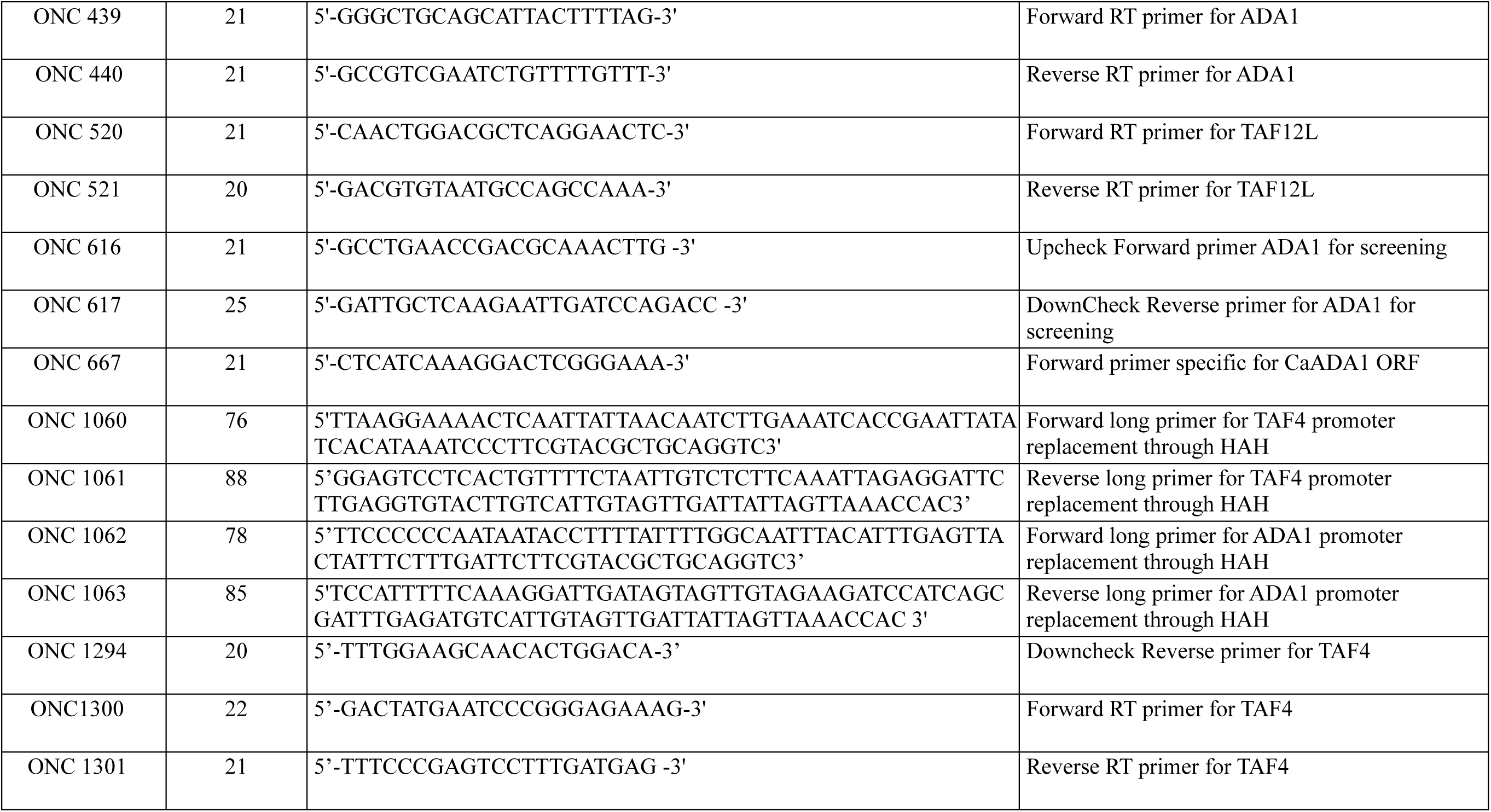
List of oligonucleotides.

**Table S3:**
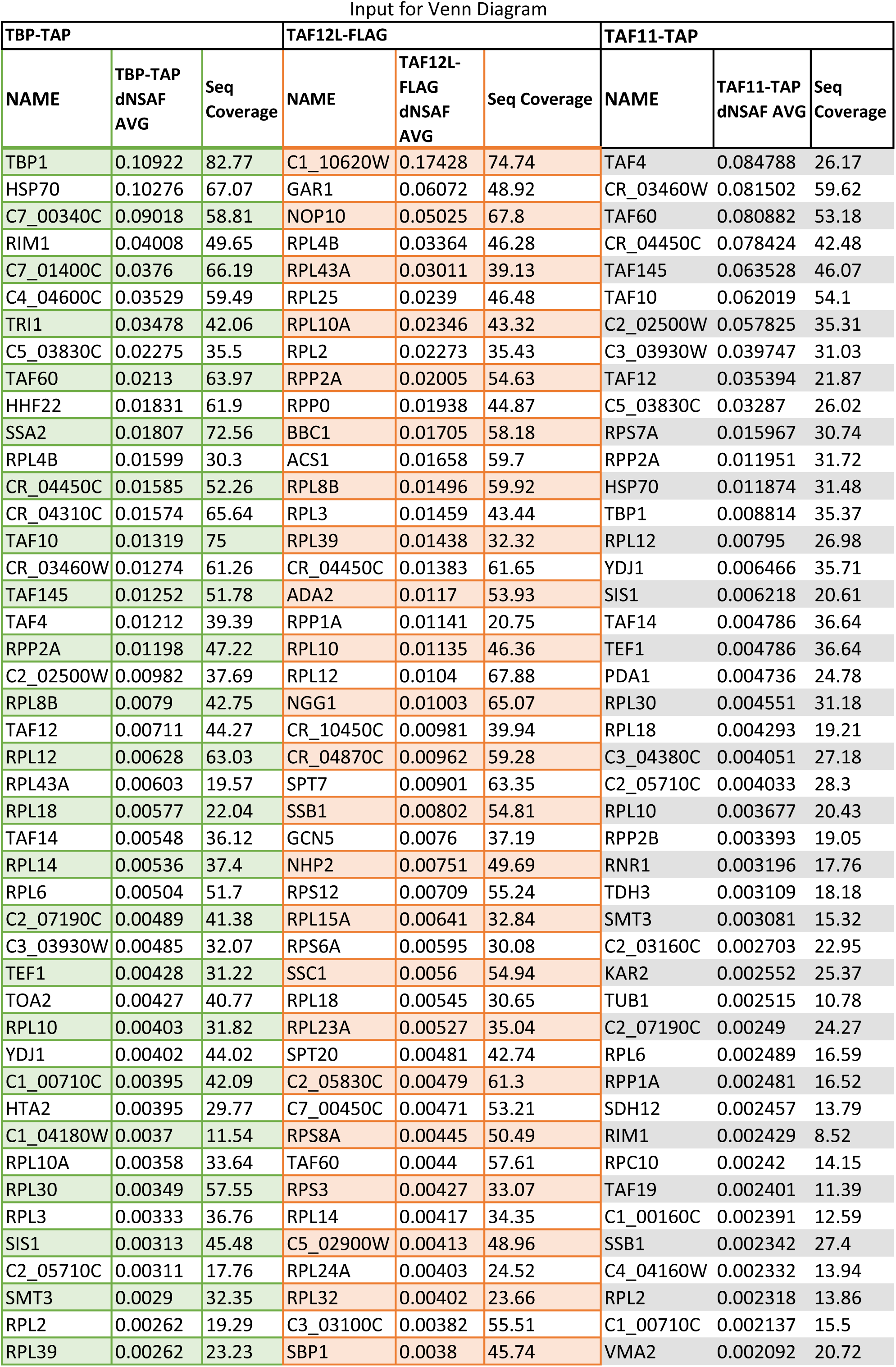
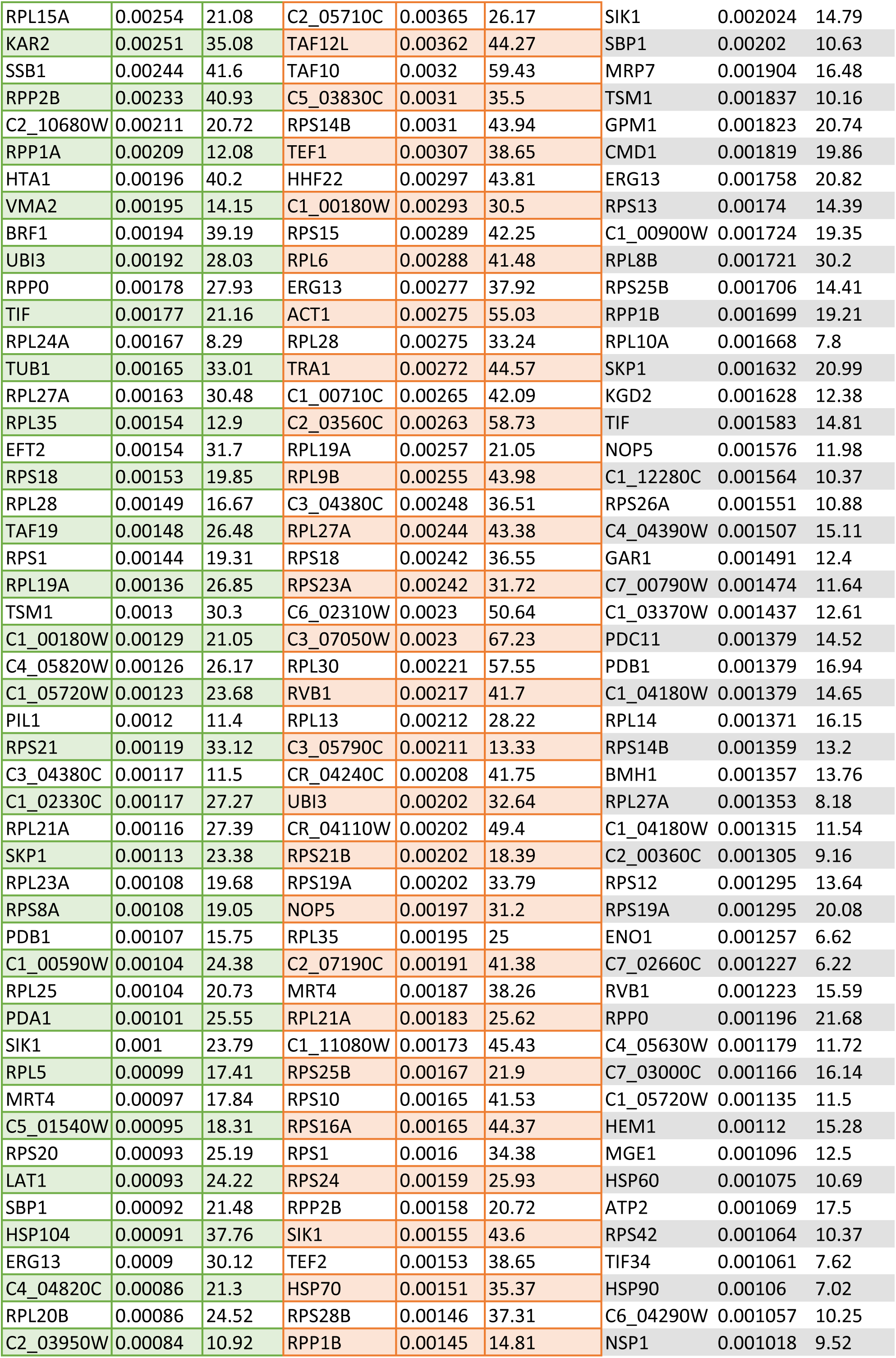
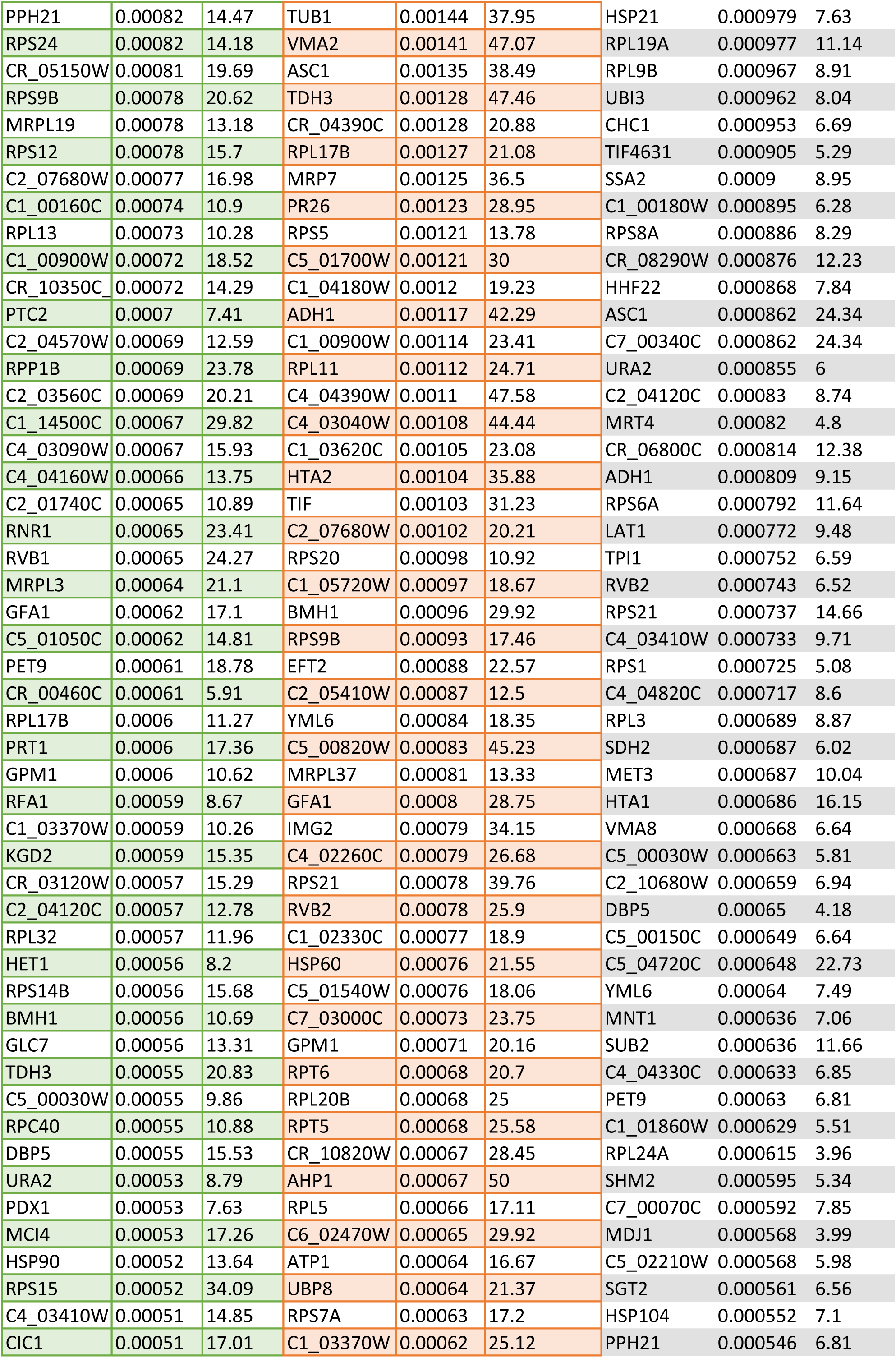
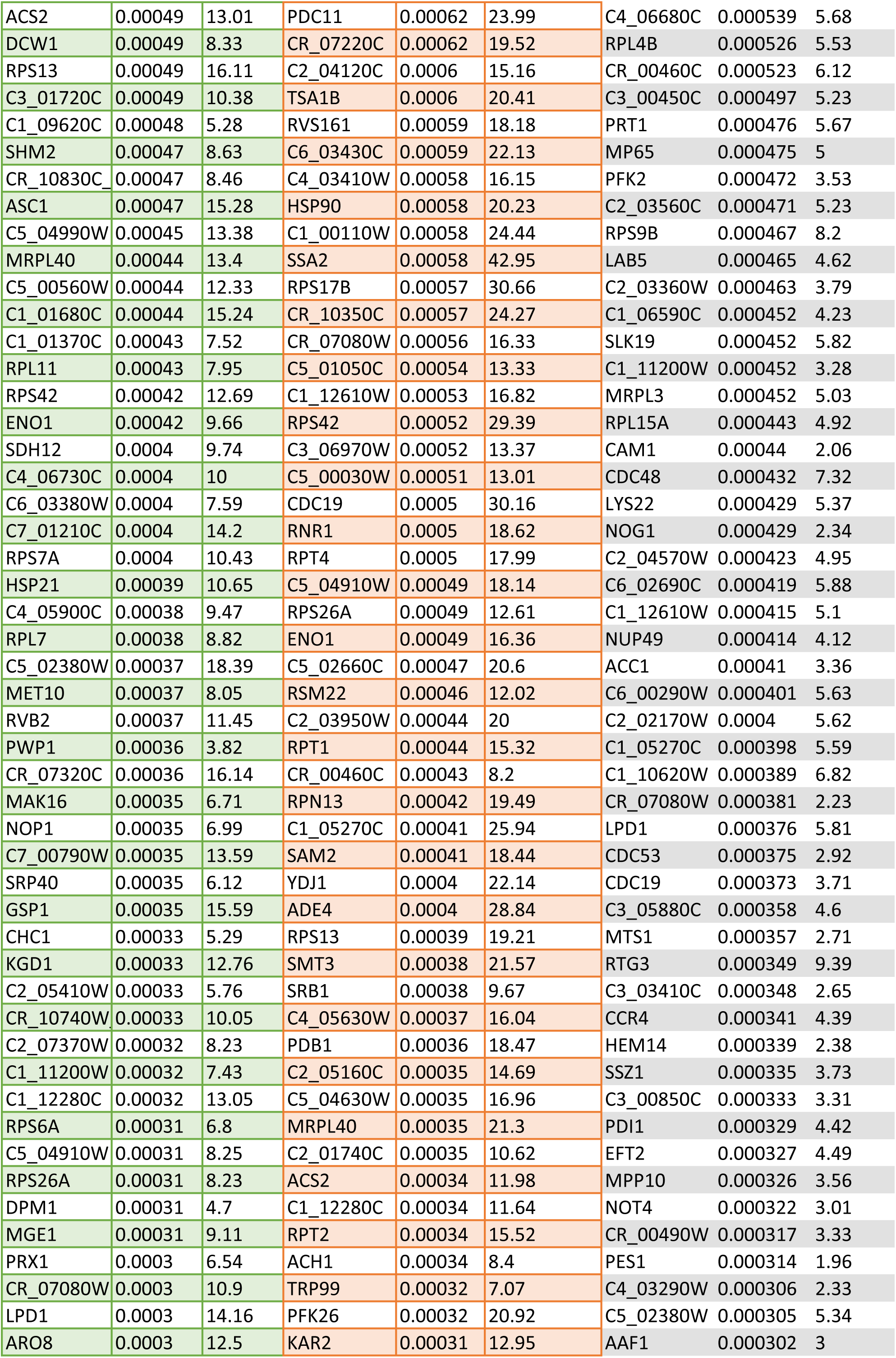
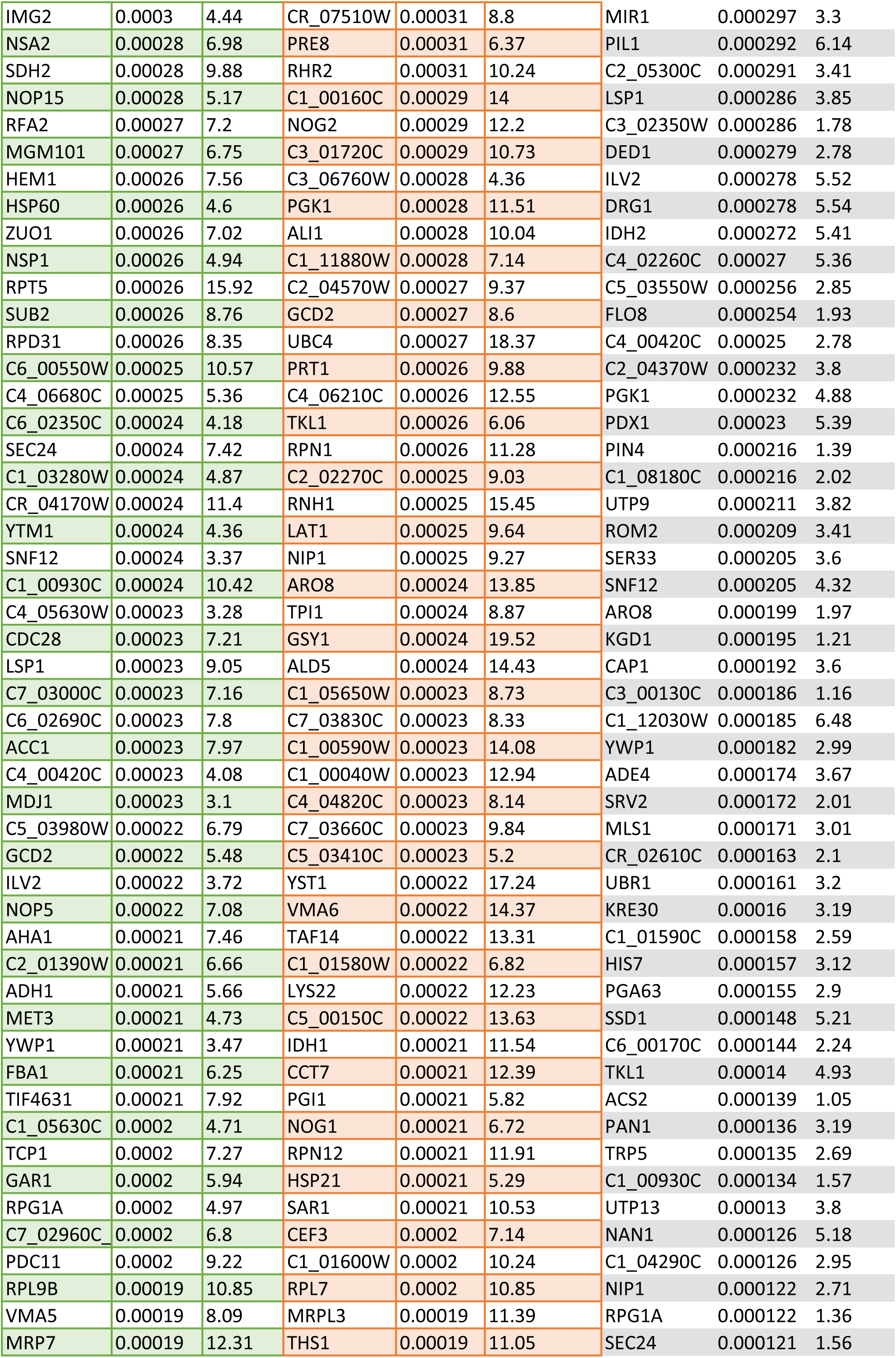
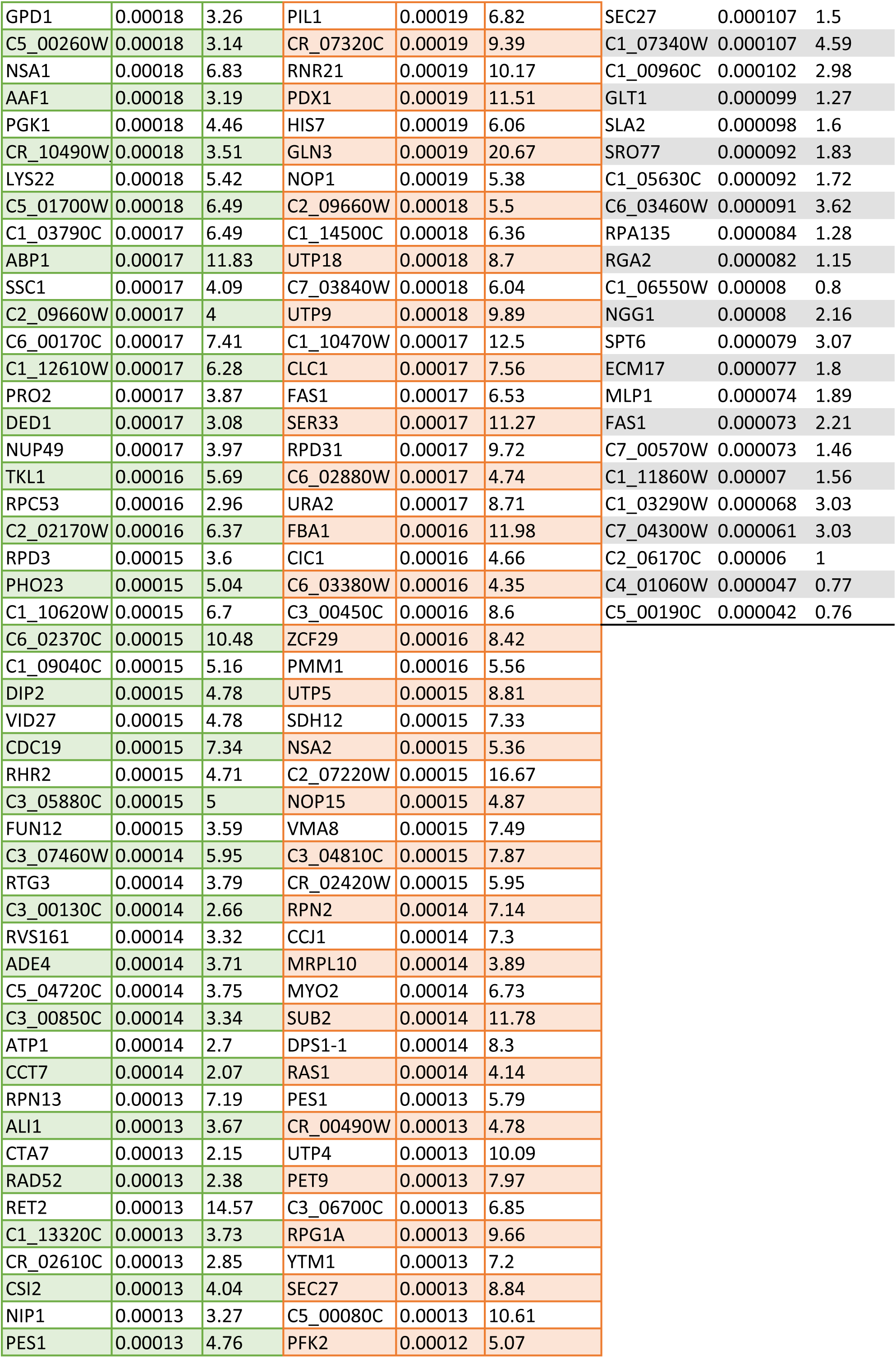
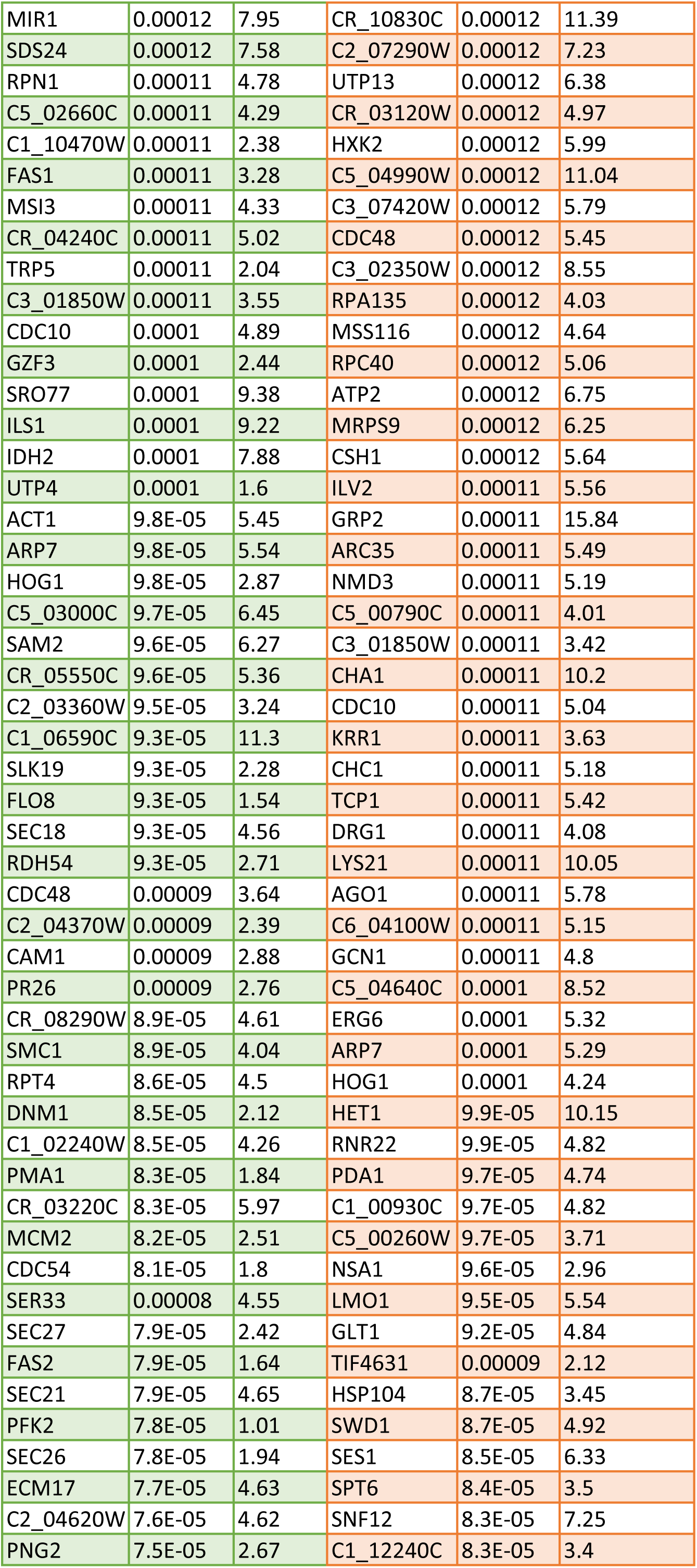
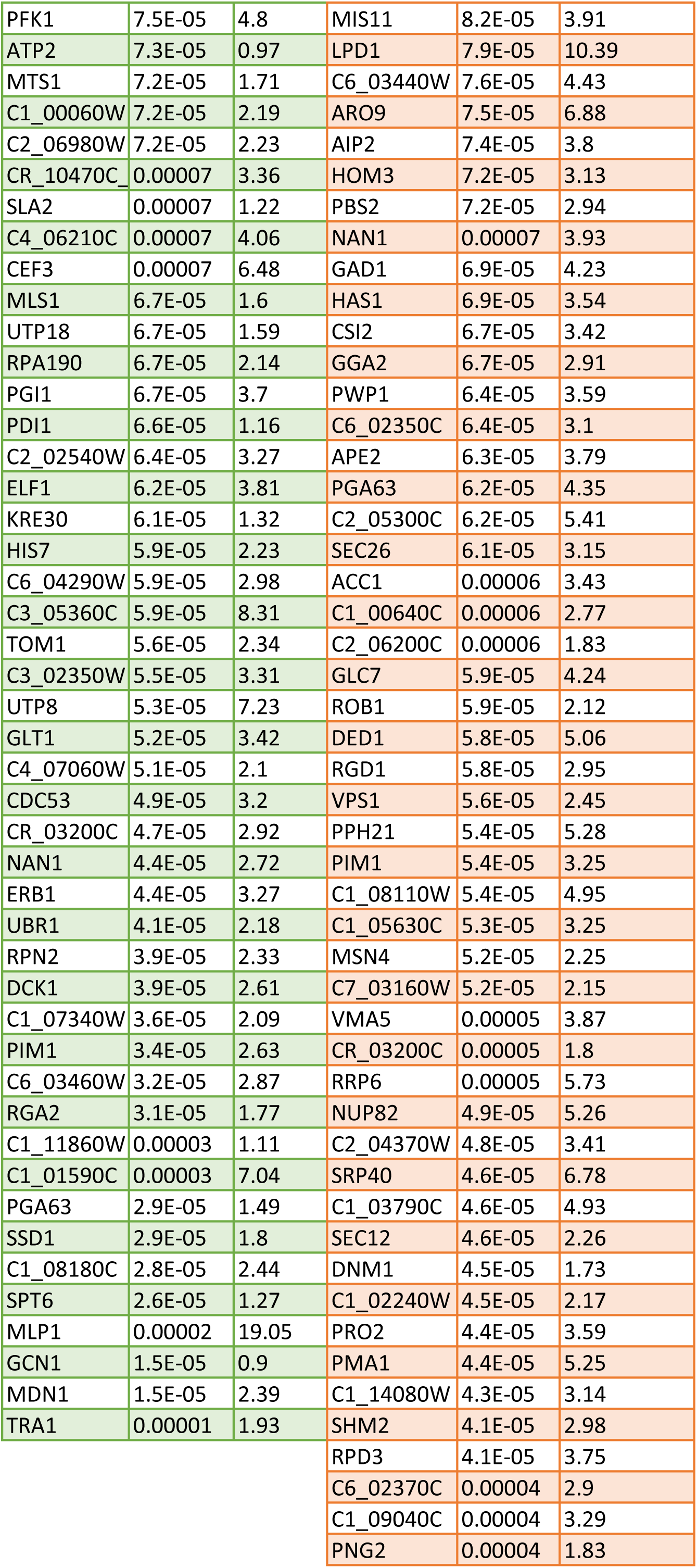
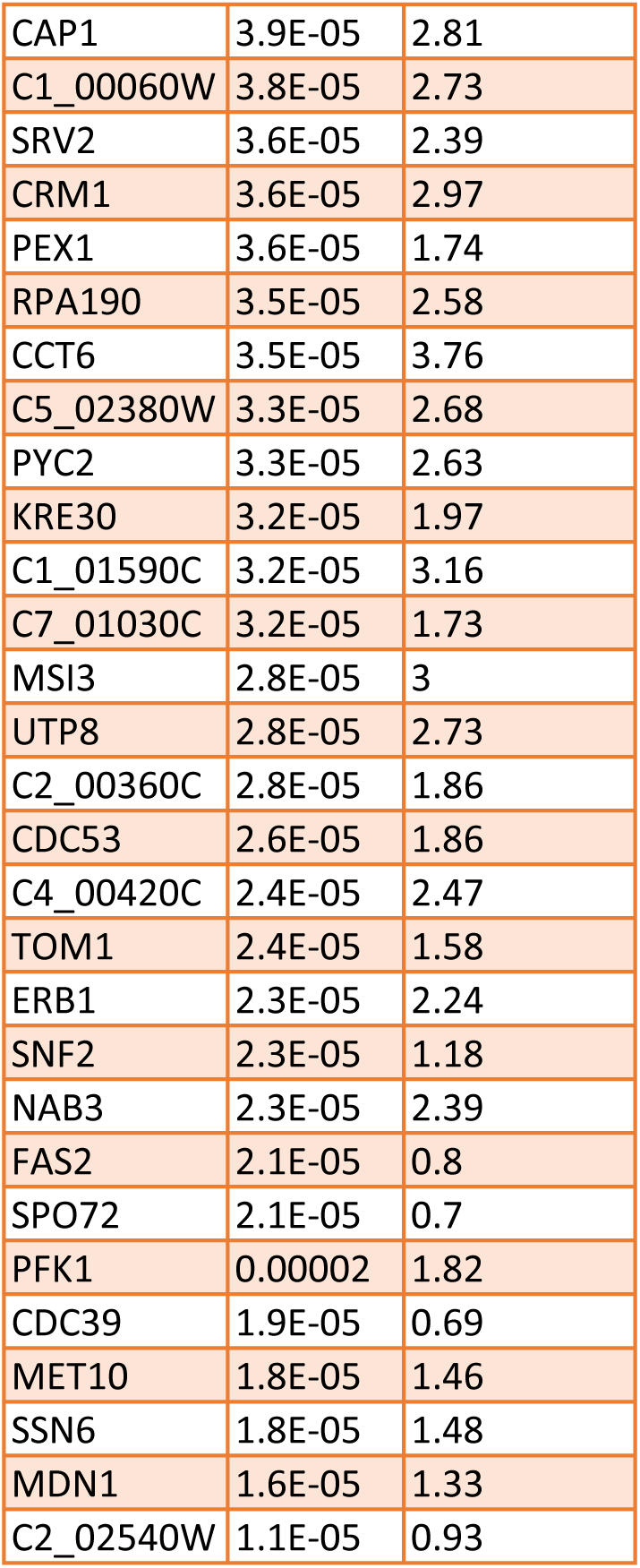
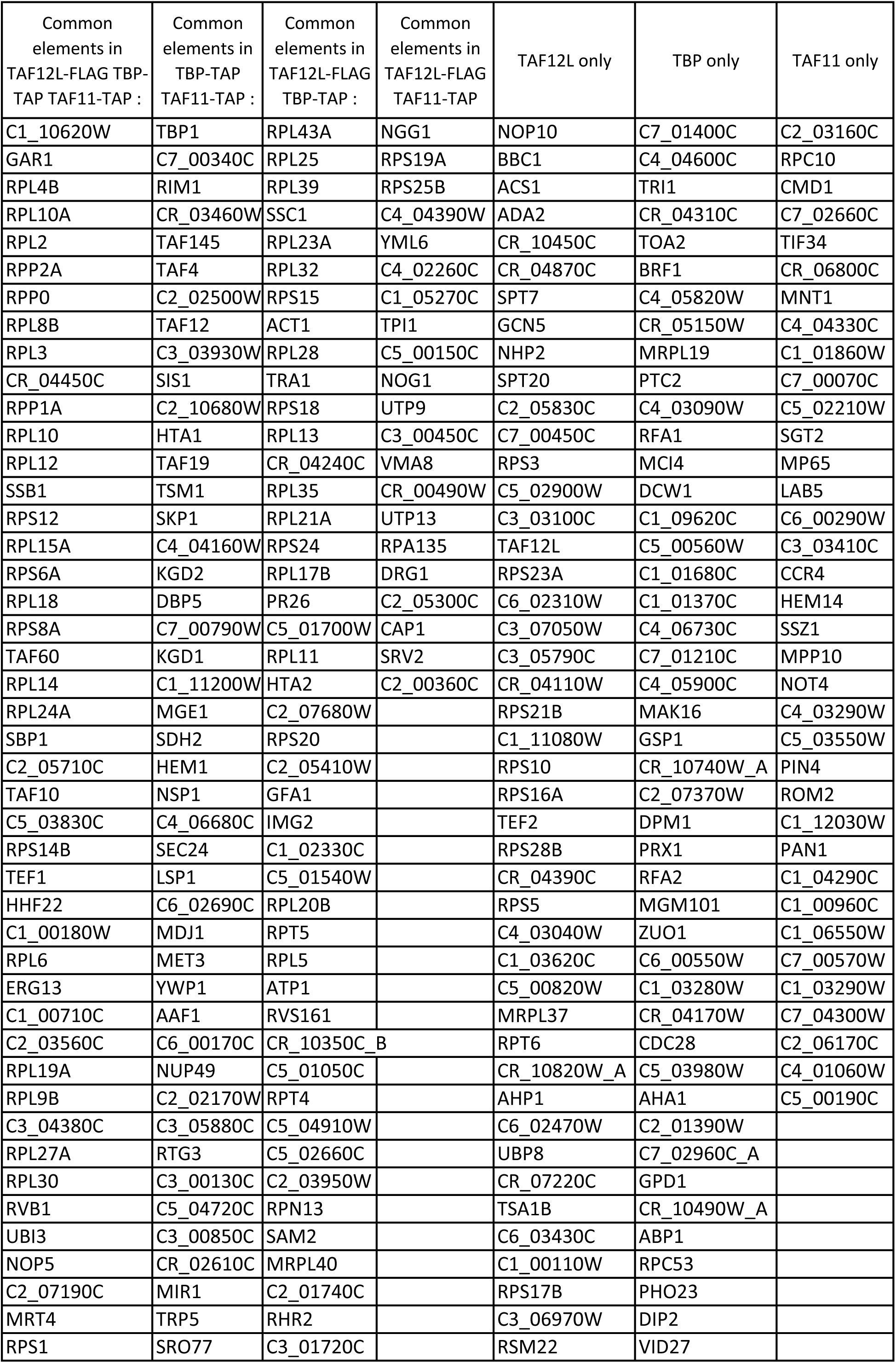
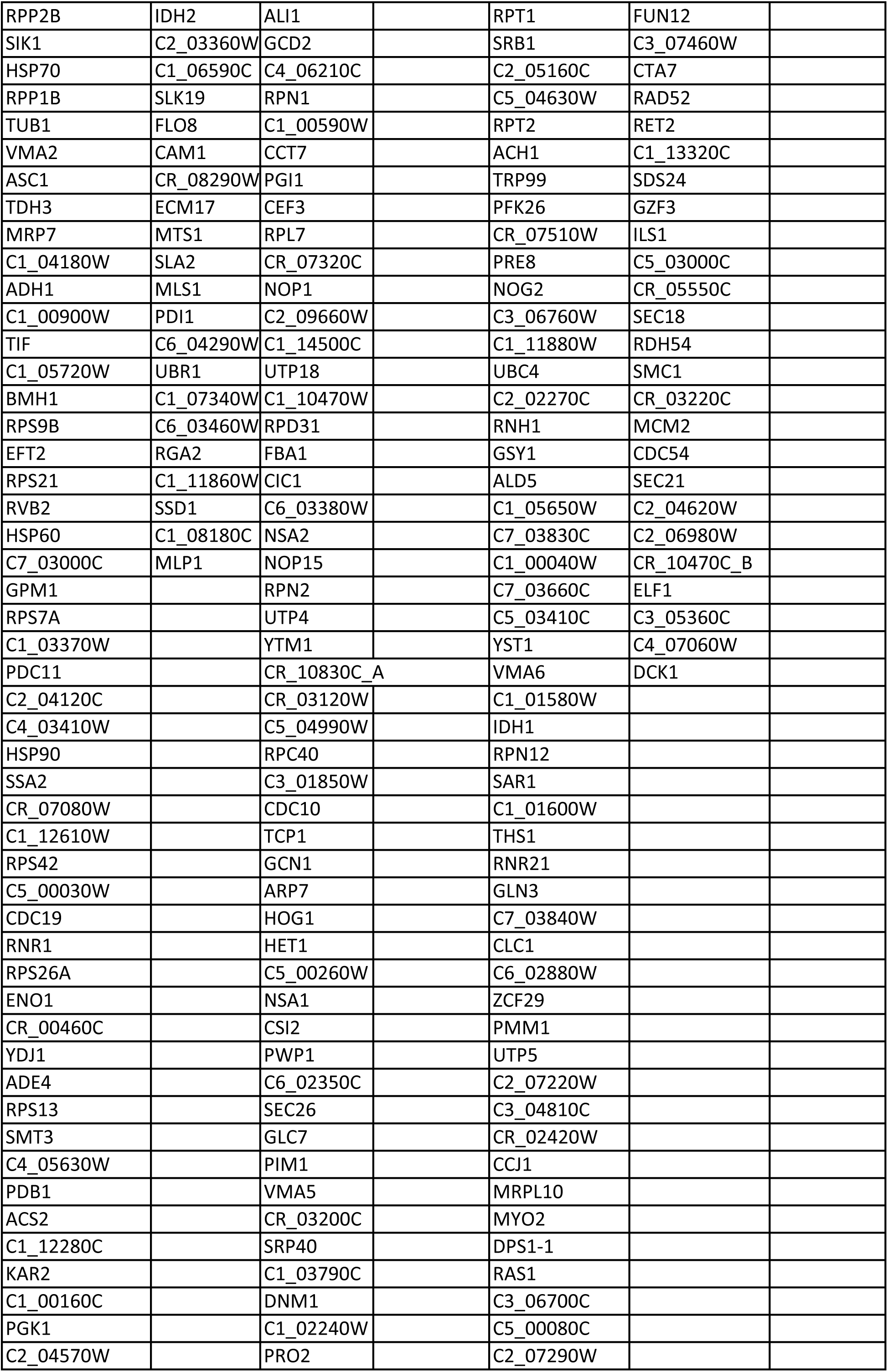
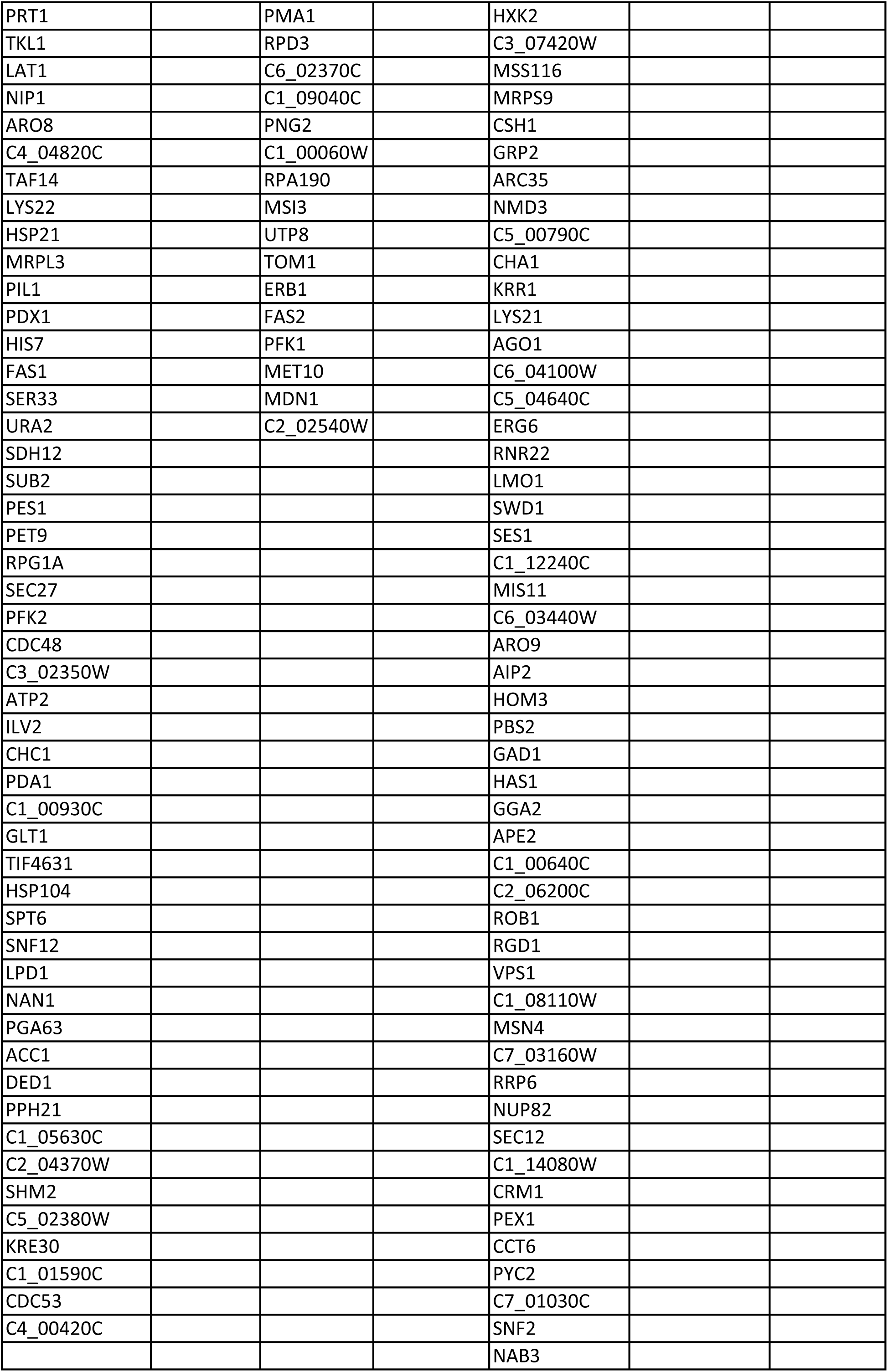
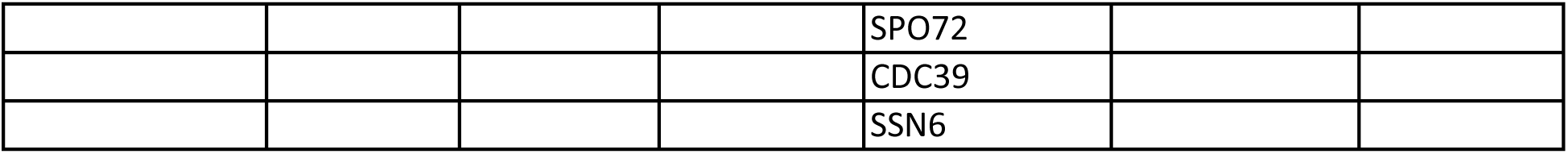
List of proteins identified by MudPIT analysis of TBP-TAP, TAF11-TAP, and TAF12L-FLAG purifications analysed by Venn diagram and the list of overlapping and non-overlapping proteins.

| TBP-TAP |  |  | TAF12L-FLAG |  |  | TAF11-TAP |  |  |
| --- | --- | --- | --- | --- | --- | --- | --- | --- |
| NAME | TBP-TAP dNSAF AVG | Seq Coverage | NAME | TAF12L-FLAG dNSAF AVG | Seq Coverage | NAME | TAF11-TAP dNSAF AVG | Seq Coverage |
| TBP1 | 0.10922 | 82.77 | C1_10620W | 0.17428 | 74.74 | TAF4 | 0.084788 | 26.17 |
| HSP70 | 0.10276 | 67.07 | GAR1 | 0.06072 | 48.92 | CR_03460W | 0.081502 | 59.62 |
| C7_00340C | 0.09018 | 58.81 | NOP10 | 0.05025 | 67.8 | TAF60 | 0.080882 | 53.18 |
| RIM1 | 0.04008 | 49.65 | RPL4B | 0.03364 | 46.28 | CR_04450C | 0.078424 | 42.48 |
| C7_01400C | 0.0376 | 66.19 | RPL43A | 0.03011 | 39.13 | TAF145 | 0.063528 | 46.07 |
| C4_04600C | 0.03529 | 59.49 | RPL25 | 0.0239 | 46.48 | TAF10 | 0.062019 | 54.1 |
| TRI1 | 0.03478 | 42.06 | RPL10A | 0.02346 | 43.32 | C2_02500W | 0.057825 | 35.31 |
| C5_03830C | 0.02275 | 35.5 | RPL2 | 0.02273 | 35.43 | C3_03930W | 0.039747 | 31.03 |
| TAF60 | 0.0213 | 63.97 | RPP2A | 0.02005 | 54.63 | TAF12 | 0.035394 | 21.87 |
| HHF22 | 0.01831 | 61.9 | RPP0 | 0.01938 | 44.87 | C5_03830C | 0.03287 | 26.02 |
| SSA2 | 0.01807 | 72.56 | BBC1 | 0.01705 | 58.18 | RPS7A | 0.015967 | 30.74 |
| RPL4B | 0.01599 | 30.3 | ACS1 | 0.01658 | 59.7 | RPP2A | 0.011951 | 31.72 |
| CR_04450C | 0.01585 | 52.26 | RPL8B | 0.01496 | 59.92 | HSP70 | 0.011874 | 31.48 |
| CR_04310C | 0.01574 | 65.64 | RPL3 | 0.01459 | 43.44 | TBP1 | 0.008814 | 35.37 |
| TAF10 | 0.01319 | 75 | RPL39 | 0.01438 | 32.32 | RPL12 | 0.00795 | 26.98 |
| CR_03460W | 0.01274 | 61.26 | CR_04450C | 0.01383 | 61.65 | YDJ1 | 0.006466 | 35.71 |
| TAF145 | 0.01252 | 51.78 | ADA2 | 0.0117 | 53.93 | SIS1 | 0.006218 | 20.61 |
| TAF4 | 0.01212 | 39.39 | RPP1A | 0.01141 | 20.75 | TAF14 | 0.004786 | 36.64 |
| RPP2A | 0.01198 | 47.22 | RPL10 | 0.01135 | 46.36 | TEF1 | 0.004786 | 36.64 |
| C2_02500W | 0.00982 | 37.69 | RPL12 | 0.0104 | 67.88 | PDA1 | 0.004736 | 24.78 |
| RPL8B | 0.0079 | 42.75 | NGG1 | 0.01003 | 65.07 | RPL30 | 0.004551 | 31.18 |
| TAF12 | 0.00711 | 44.27 | CR_10450C | 0.00981 | 39.94 | RPL18 | 0.004293 | 19.21 |
| RPL12 | 0.00628 | 63.03 | CR_04870C | 0.00962 | 59.28 | C3_04380C | 0.004051 | 27.18 |
| RPL43A | 0.00603 | 19.57 | SPT7 | 0.00901 | 63.35 | C2_05710C | 0.004033 | 28.3 |
| RPL18 | 0.00577 | 22.04 | SSB1 | 0.00802 | 54.81 | RPL10 | 0.003677 | 20.43 |
| TAF14 | 0.00548 | 36.12 | GCN5 | 0.0076 | 37.19 | RPP2B | 0.003393 | 19.05 |
| RPL14 | 0.00536 | 37.4 | NHP2 | 0.00751 | 49.69 | RNR1 | 0.003196 | 17.76 |
| RPL6 | 0.00504 | 51.7 | RPS12 | 0.00709 | 55.24 | TDH3 | 0.003109 | 18.18 |
| C2_07190C | 0.00489 | 41.38 | RPL15A | 0.00641 | 32.84 | SMT3 | 0.003081 | 15.32 |
| C3_03930W | 0.00485 | 32.07 | RPS6A | 0.00595 | 30.08 | C2_03160C | 0.002703 | 22.95 |
| TEF1 | 0.00428 | 31.22 | SSC1 | 0.0056 | 54.94 | KAR2 | 0.002552 | 25.37 |
| TOA2 | 0.00427 | 40.77 | RPL18 | 0.00545 | 30.65 | TUB1 | 0.002515 | 10.78 |
| RPL10 | 0.00403 | 31.82 | RPL23A | 0.00527 | 35.04 | C2_07190C | 0.00249 | 24.27 |
| YDJ1 | 0.00402 | 44.02 | SPT20 | 0.00481 | 42.74 | RPL6 | 0.002489 | 16.59 |
| C1_00710C | 0.00395 | 42.09 | C2_05830C | 0.00479 | 61.3 | RPP1A | 0.002481 | 16.52 |
| HTA2 | 0.00395 | 29.77 | C7_00450C | 0.00471 | 53.21 | SDH12 | 0.002457 | 13.79 |
| C1_04180W | 0.0037 | 11.54 | RPS8A | 0.00445 | 50.49 | RIM1 | 0.002429 | 8.52 |
| RPL10A | 0.00358 | 33.64 | TAF60 | 0.0044 | 57.61 | RPC10 | 0.00242 | 14.15 |
| RPL30 | 0.00349 | 57.55 | RPS3 | 0.00427 | 33.07 | TAF19 | 0.002401 | 11.39 |
| RPL3 | 0.00333 | 36.76 | RPL14 | 0.00417 | 34.35 | C1_00160C | 0.002391 | 12.59 |
| SIS1 | 0.00313 | 45.48 | C5_02900W | 0.00413 | 48.96 | SSB1 | 0.002342 | 27.4 |
| C2_05710C | 0.00311 | 17.76 | RPL24A | 0.00403 | 24.52 | C4_04160W | 0.002332 | 13.94 |
| SMT3 | 0.0029 | 32.35 | RPL32 | 0.00402 | 23.66 | RPL2 | 0.002318 | 13.86 |
| RPL2 | 0.00262 | 19.29 | C3_03100C | 0.00382 | 55.51 | C1_00710C | 0.002137 | 15.5 |
| RPL39 | 0.00262 | 23.23 | SBP1 | 0.0038 | 45.74 | VMA2 | 0.002092 | 20.72 |
| RPL15A | 0.00254 | 21.08 | C2_05710C | 0.00365 | 26.17 | SIK1 | 0.002024 | 14.79 |
| KAR2 | 0.00251 | 35.08 | TAF12L | 0.00362 | 44.27 | SBP1 | 0.00202 | 10.63 |
| SSB1 | 0.00244 | 41.6 | TAF10 | 0.0032 | 59.43 | MRP7 | 0.001904 | 16.48 |
| RPP2B | 0.00233 | 40.93 | C5_03830C | 0.0031 | 35.5 | TSM1 | 0.001837 | 10.16 |
| C2_10680W | 0.00211 | 20.72 | RPS14B | 0.0031 | 43.94 | GPM1 | 0.001823 | 20.74 |
| RPP1A | 0.00209 | 12.08 | TEF1 | 0.00307 | 38.65 | CMD1 | 0.001819 | 19.86 |
| HTA1 | 0.00196 | 40.2 | HHF22 | 0.00297 | 43.81 | ERG13 | 0.001758 | 20.82 |
| VMA2 | 0.00195 | 14.15 | C1_00180W | 0.00293 | 30.5 | RPS13 | 0.00174 | 14.39 |
| BRF1 | 0.00194 | 39.19 | RPS15 | 0.00289 | 42.25 | C1_00900W | 0.001724 | 19.35 |
| UBI3 | 0.00192 | 28.03 | RPL6 | 0.00288 | 41.48 | RPL8B | 0.001721 | 30.2 |
| RPP0 | 0.00178 | 27.93 | ERG13 | 0.00277 | 37.92 | RPS25B | 0.001706 | 14.41 |
| TIF | 0.00177 | 21.16 | ACT1 | 0.00275 | 55.03 | RPP1B | 0.001699 | 19.21 |
| RPL24A | 0.00167 | 8.29 | RPL28 | 0.00275 | 33.24 | RPL10A | 0.001668 | 7.8 |
| TUB1 | 0.00165 | 33.01 | TRA1 | 0.00272 | 44.57 | SKP1 | 0.001632 | 20.99 |
| RPL27A | 0.00163 | 30.48 | C1_00710C | 0.00265 | 42.09 | KGD2 | 0.001628 | 12.38 |
| RPL35 | 0.00154 | 12.9 | C2_03560C | 0.00263 | 58.73 | TIF | 0.001583 | 14.81 |
| EFT2 | 0.00154 | 31.7 | RPL19A | 0.00257 | 21.05 | NOP5 | 0.001576 | 11.98 |
| RPS18 | 0.00153 | 19.85 | RPL9B | 0.00255 | 43.98 | C1_12280C | 0.001564 | 10.37 |
| RPL28 | 0.00149 | 16.67 | C3_04380C | 0.00248 | 36.51 | RPS26A | 0.001551 | 10.88 |
| TAF19 | 0.00148 | 26.48 | RPL27A | 0.00244 | 43.38 | C4_04390W | 0.001507 | 15.11 |
| RPS1 | 0.00144 | 19.31 | RPS18 | 0.00242 | 36.55 | GAR1 | 0.001491 | 12.4 |
| RPL19A | 0.00136 | 26.85 | RPS23A | 0.00242 | 31.72 | C7_00790W | 0.001474 | 11.64 |
| TSM1 | 0.0013 | 30.3 | C6_02310W | 0.0023 | 50.64 | C1_03370W | 0.001437 | 12.61 |
| C1_00180W | 0.00129 | 21.05 | C3_07050W | 0.0023 | 67.23 | PDC11 | 0.001379 | 14.52 |
| C4_05820W | 0.00126 | 26.17 | RPL30 | 0.00221 | 57.55 | PDB1 | 0.001379 | 16.94 |
| C1_05720W | 0.00123 | 23.68 | RVB1 | 0.00217 | 41.7 | C1_04180W | 0.001379 | 14.65 |
| PIL1 | 0.0012 | 11.4 | RPL13 | 0.00212 | 28.22 | RPL14 | 0.001371 | 16.15 |
| RPS21 | 0.00119 | 33.12 | C3_05790C | 0.00211 | 13.33 | RPS14B | 0.001359 | 13.2 |
| C3_04380C | 0.00117 | 11.5 | CR_04240C | 0.00208 | 41.75 | BMH1 | 0.001357 | 13.76 |
| C1_02330C | 0.00117 | 27.27 | UBI3 | 0.00202 | 32.64 | RPL27A | 0.001353 | 8.18 |
| RPL21A | 0.00116 | 27.39 | CR_04110W | 0.00202 | 49.4 | C1_04180W | 0.001315 | 11.54 |
| SKP1 | 0.00113 | 23.38 | RPS21B | 0.00202 | 18.39 | C2_00360C | 0.001305 | 9.16 |
| RPL23A | 0.00108 | 19.68 | RPS19A | 0.00202 | 33.79 | RPS12 | 0.001295 | 13.64 |
| RPS8A | 0.00108 | 19.05 | NOP5 | 0.00197 | 31.2 | RPS19A | 0.001295 | 20.08 |
| PDB1 | 0.00107 | 15.75 | RPL35 | 0.00195 | 25 | ENO1 | 0.001257 | 6.62 |
| C1_00590W | 0.00104 | 24.38 | C2_07190C | 0.00191 | 41.38 | C7_02660C | 0.001227 | 6.22 |
| RPL25 | 0.00104 | 20.73 | MRT4 | 0.00187 | 38.26 | RVB1 | 0.001223 | 15.59 |
| PDA1 | 0.00101 | 25.55 | RPL21A | 0.00183 | 25.62 | RPP0 | 0.001196 | 21.68 |
| SIK1 | 0.001 | 23.79 | C1_11080W | 0.00173 | 45.43 | C4_05630W | 0.001179 | 11.72 |
| RPL5 | 0.00099 | 17.41 | RPS25B | 0.00167 | 21.9 | C7_03000C | 0.001166 | 16.14 |
| MRT4 | 0.00097 | 17.84 | RPS10 | 0.00165 | 41.53 | C1_05720W | 0.001135 | 11.5 |
| C5_01540W | 0.00095 | 18.31 | RPS16A | 0.00165 | 44.37 | HEM1 | 0.00112 | 15.28 |
| RPS20 | 0.00093 | 25.19 | RPS1 | 0.0016 | 34.38 | MGE1 | 0.001096 | 12.5 |
| LAT1 | 0.00093 | 24.22 | RPS24 | 0.00159 | 25.93 | HSP60 | 0.001075 | 10.69 |
| SBP1 | 0.00092 | 21.48 | RPP2B | 0.00158 | 20.72 | ATP2 | 0.001069 | 17.5 |
| HSP104 | 0.00091 | 37.76 | SIK1 | 0.00155 | 43.6 | RPS42 | 0.001064 | 10.37 |
| ERG13 | 0.0009 | 30.12 | TEF2 | 0.00153 | 38.65 | TIF34 | 0.001061 | 7.62 |
| C4_04820C | 0.00086 | 21.3 | HSP70 | 0.00151 | 35.37 | HSP90 | 0.00106 | 7.02 |
| RPL20B | 0.00086 | 24.52 | RPS28B | 0.00146 | 37.31 | C6_04290W | 0.001057 | 10.25 |
| C2_03950W | 0.00084 | 10.92 | RPP1B | 0.00145 | 14.81 | NSP1 | 0.001018 | 9.52 |
| PPH21 | 0.00082 | 14.47 | TUB1 | 0.00144 | 37.95 | HSP21 | 0.000979 | 7.63 |
| RPS24 | 0.00082 | 14.18 | VMA2 | 0.00141 | 47.07 | RPL19A | 0.000977 | 11.14 |
| CR_05150W | 0.00081 | 19.69 | ASC1 | 0.00135 | 38.49 | RPL9B | 0.000967 | 8.91 |
| RPS9B | 0.00078 | 20.62 | TDH3 | 0.00128 | 47.46 | UBI3 | 0.000962 | 8.04 |
| MRPL19 | 0.00078 | 13.18 | CR_04390C | 0.00128 | 20.88 | CHC1 | 0.000953 | 6.69 |
| RPS12 | 0.00078 | 15.7 | RPL17B | 0.00127 | 21.08 | TIF4631 | 0.000905 | 5.29 |
| C2_07680W | 0.00077 | 16.98 | MRP7 | 0.00125 | 36.5 | SSA2 | 0.0009 | 8.95 |
| C1_00160C | 0.00074 | 10.9 | PR26 | 0.00123 | 28.95 | C1_00180W | 0.000895 | 6.28 |
| RPL13 | 0.00073 | 10.28 | RPS5 | 0.00121 | 13.78 | RPS8A | 0.000886 | 8.29 |
| C1_00900W | 0.00072 | 18.52 | C5_01700W | 0.00121 | 30 | CR_08290W | 0.000876 | 12.23 |
| CR_10350C | 0.00072 | 14.29 | C1_04180W | 0.0012 | 19.23 | HHF22 | 0.000868 | 7.84 |
| PTC2 | 0.0007 | 7.41 | ADH1 | 0.00117 | 42.29 | ASC1 | 0.000862 | 24.34 |
| C2_04570W | 0.00069 | 12.59 | C1_00900W | 0.00114 | 23.41 | C7_00340C | 0.000862 | 24.34 |
| RPP1B | 0.00069 | 23.78 | RPL11 | 0.00112 | 24.71 | URA2 | 0.000855 | 6 |
| C2_03560C | 0.00069 | 20.21 | C4_04390W | 0.0011 | 47.58 | C2_04120C | 0.00083 | 8.74 |
| C1_14500C | 0.00067 | 29.82 | C4_03040W | 0.00108 | 44.44 | MRT4 | 0.00082 | 4.8 |
| C4_03090W | 0.00067 | 15.93 | C1_03620C | 0.00105 | 23.08 | CR_06800C | 0.000814 | 12.38 |
| C4_04160W | 0.00066 | 13.75 | HTA2 | 0.00104 | 35.88 | ADH1 | 0.000809 | 9.15 |
| C2_01740C | 0.00065 | 10.89 | TIF | 0.00103 | 31.23 | RPS6A | 0.000792 | 11.64 |
| RNR1 | 0.00065 | 23.41 | C2_07680W | 0.00102 | 20.21 | LAT1 | 0.000772 | 9.48 |
| RVB1 | 0.00065 | 24.27 | RPS20 | 0.00098 | 10.92 | TPI1 | 0.000752 | 6.59 |
| MRPL3 | 0.00064 | 21.1 | C1_05720W | 0.00097 | 18.67 | RVB2 | 0.000743 | 6.52 |
| GFA1 | 0.00062 | 17.1 | BMH1 | 0.00096 | 29.92 | RPS21 | 0.000737 | 14.66 |
| C5_01050C | 0.00062 | 14.81 | RPS9B | 0.00093 | 17.46 | C4_03410W | 0.000733 | 9.71 |
| PET9 | 0.00061 | 18.78 | EFT2 | 0.00088 | 22.57 | RPS1 | 0.000725 | 5.08 |
| CR_00460C | 0.00061 | 5.91 | C2_05410W | 0.00087 | 12.5 | C4_04820C | 0.000717 | 8.6 |
| RPL17B | 0.0006 | 11.27 | YML6 | 0.00084 | 18.35 | RPL3 | 0.000689 | 8.87 |
| PRT1 | 0.0006 | 17.36 | C5_00820W | 0.00083 | 45.23 | SDH2 | 0.000687 | 6.02 |
| GPM1 | 0.0006 | 10.62 | MRPL37 | 0.00081 | 13.33 | MET3 | 0.000687 | 10.04 |
| RFA1 | 0.00059 | 8.67 | GFA1 | 0.0008 | 28.75 | HTA1 | 0.000686 | 16.15 |
| C1_03370W | 0.00059 | 10.26 | IMG2 | 0.00079 | 34.15 | VMA8 | 0.000668 | 6.64 |
| KGD2 | 0.00059 | 15.35 | C4_02260C | 0.00079 | 26.68 | C5_00030W | 0.000663 | 5.81 |
| CR_03120W | 0.00057 | 15.29 | RPS21 | 0.00078 | 39.76 | C2_10680W | 0.000659 | 6.94 |
| C2_04120C | 0.00057 | 12.78 | RVB2 | 0.00078 | 25.9 | DBP5 | 0.00065 | 4.18 |
| RPL32 | 0.00057 | 11.96 | C1_02330C | 0.00077 | 18.9 | C5_00150C | 0.000649 | 6.64 |
| HET1 | 0.00056 | 8.2 | HSP60 | 0.00076 | 21.55 | C5_04720C | 0.000648 | 22.73 |
| RPS14B | 0.00056 | 15.68 | C5_01540W | 0.00076 | 18.06 | YML6 | 0.00064 | 7.49 |
| BMH1 | 0.00056 | 10.69 | C7_03000C | 0.00073 | 23.75 | MNT1 | 0.000636 | 7.06 |
| GLC7 | 0.00056 | 13.31 | GPM1 | 0.00071 | 20.16 | SUB2 | 0.000636 | 11.66 |
| TDH3 | 0.00055 | 20.83 | RPT6 | 0.00068 | 20.7 | C4_04330C | 0.000633 | 6.85 |
| C5_00030W | 0.00055 | 9.86 | RPL20B | 0.00068 | 25 | PET9 | 0.00063 | 6.81 |
| RPC40 | 0.00055 | 10.88 | RPT5 | 0.00068 | 25.58 | C1_01860W | 0.000629 | 5.51 |
| DBP5 | 0.00055 | 15.53 | CR_10820W | 0.00067 | 28.45 | RPL24A | 0.000615 | 3.96 |
| URA2 | 0.00053 | 8.79 | AHP1 | 0.00067 | 50 | SHM2 | 0.000595 | 5.34 |
| PDX1 | 0.00053 | 7.63 | RPL5 | 0.00066 | 17.11 | C7_00070C | 0.000592 | 7.85 |
| MCI4 | 0.00053 | 17.26 | C6_02470W | 0.00065 | 29.92 | MDJ1 | 0.000568 | 3.99 |
| HSP90 | 0.00052 | 13.64 | ATP1 | 0.00064 | 16.67 | C5_02210W | 0.000568 | 5.98 |
| RPS15 | 0.00052 | 34.09 | UBP8 | 0.00064 | 21.37 | SGT2 | 0.000561 | 6.56 |
| C4_03410W | 0.00051 | 14.85 | RPS7A | 0.00063 | 17.2 | HSP104 | 0.000552 | 7.1 |
| CIC1 | 0.00051 | 17.01 | C1_03370W | 0.00062 | 25.12 | PPH21 | 0.000546 | 6.81 |
| ACS2 | 0.00049 | 13.01 | PDC11 | 0.00062 | 23.99 | C4_06680C | 0.000539 | 5.68 |
| DCW1 | 0.00049 | 8.33 | CR_07220C | 0.00062 | 19.52 | RPL4B | 0.000526 | 5.53 |
| RPS13 | 0.00049 | 16.11 | C2_04120C | 0.0006 | 15.16 | CR_00460C | 0.000523 | 6.12 |
| C3_01720C | 0.00049 | 10.38 | TSA1B | 0.0006 | 20.41 | C3_00450C | 0.000497 | 5.23 |
| C1_09620C | 0.00048 | 5.28 | RVS161 | 0.00059 | 18.18 | PRT1 | 0.000476 | 5.67 |
| SHM2 | 0.00047 | 8.63 | C6_03430C | 0.00059 | 22.13 | MP65 | 0.000475 | 5 |
| CR_10830C | 0.00047 | 8.46 | C4_03410W | 0.00058 | 16.15 | PFK2 | 0.000472 | 3.53 |
| ASC1 | 0.00047 | 15.28 | HSP90 | 0.00058 | 20.23 | C2_03560C | 0.000471 | 5.23 |
| C5_04990W | 0.00045 | 13.38 | C1_00110W | 0.00058 | 24.44 | RPS9B | 0.000467 | 8.2 |
| MRPL40 | 0.00044 | 13.4 | SSA2 | 0.00058 | 42.95 | LAB5 | 0.000465 | 4.62 |
| C5_00560W | 0.00044 | 12.33 | RPS17B | 0.00057 | 30.66 | C2_03360W | 0.000463 | 3.79 |
| C1_01680C | 0.00044 | 15.24 | CR_10350C | 0.00057 | 24.27 | C1_06590C | 0.000452 | 4.23 |
| C1_01370C | 0.00043 | 7.52 | CR_07080W | 0.00056 | 16.33 | SLK19 | 0.000452 | 5.82 |
| RPL11 | 0.00043 | 7.95 | C5_01050C | 0.00054 | 13.33 | C1_11200W | 0.000452 | 3.28 |
| RPS42 | 0.00042 | 12.69 | C1_12610W | 0.00053 | 16.82 | MRPL3 | 0.000452 | 5.03 |
| ENO1 | 0.00042 | 9.66 | RPS42 | 0.00052 | 29.39 | RPL15A | 0.000443 | 4.92 |
| SDH12 | 0.0004 | 9.74 | C3_06970W | 0.00052 | 13.37 | CAM1 | 0.00044 | 2.06 |
| C4_06730C | 0.0004 | 10 | C5_00030W | 0.00051 | 13.01 | CDC48 | 0.000432 | 7.32 |
| C6_03380W | 0.0004 | 7.59 | CDC19 | 0.0005 | 30.16 | LYS22 | 0.000429 | 5.37 |
| C7_01210C | 0.0004 | 14.2 | RNR1 | 0.0005 | 18.62 | NOG1 | 0.000429 | 2.34 |
| RPS7A | 0.0004 | 10.43 | RPT4 | 0.0005 | 17.99 | C2_04570W | 0.000423 | 4.95 |
| HSP21 | 0.00039 | 10.65 | C5_04910W | 0.00049 | 18.14 | C6_02690C | 0.000419 | 5.88 |
| C4_05900C | 0.00038 | 9.47 | RPS26A | 0.00049 | 12.61 | C1_12610W | 0.000415 | 5.1 |
| RPL7 | 0.00038 | 8.82 | ENO1 | 0.00049 | 16.36 | NUP49 | 0.000414 | 4.12 |
| C5_02380W | 0.00037 | 18.39 | C5_02660C | 0.00047 | 20.6 | ACC1 | 0.00041 | 3.36 |
| MET10 | 0.00037 | 8.05 | RSM22 | 0.00046 | 12.02 | C6_00290W | 0.000401 | 5.63 |
| RVB2 | 0.00037 | 11.45 | C2_03950W | 0.00044 | 20 | C2_02170W | 0.0004 | 5.62 |
| PWP1 | 0.00036 | 3.82 | RPT1 | 0.00044 | 15.32 | C1_05270C | 0.000398 | 5.59 |
| CR_07320C | 0.00036 | 16.14 | CR_00460C | 0.00043 | 8.2 | C1_10620W | 0.000389 | 6.82 |
| MAK16 | 0.00035 | 6.71 | RPN13 | 0.00042 | 19.49 | CR_07080W | 0.000381 | 2.23 |
| NOP1 | 0.00035 | 6.99 | C1_05270C | 0.00041 | 25.94 | LPD1 | 0.000376 | 5.81 |
| C7_00790W | 0.00035 | 13.59 | SAM2 | 0.00041 | 18.44 | CDC53 | 0.000375 | 2.92 |
| SRP40 | 0.00035 | 6.12 | YDJ1 | 0.0004 | 22.14 | CDC19 | 0.000373 | 3.71 |
| GSP1 | 0.00035 | 15.59 | ADE4 | 0.0004 | 28.84 | C3_05880C | 0.000358 | 4.6 |
| CHC1 | 0.00033 | 5.29 | RPS13 | 0.00039 | 19.21 | MTS1 | 0.000357 | 2.71 |
| KGD1 | 0.00033 | 12.76 | SMT3 | 0.00038 | 21.57 | RTG3 | 0.000349 | 9.39 |
| C2_05410W | 0.00033 | 5.76 | SRB1 | 0.00038 | 9.67 | C3_03410C | 0.000348 | 2.65 |
| CR_10740W | 0.00033 | 10.05 | C4_05630W | 0.00037 | 16.04 | CCR4 | 0.000341 | 4.39 |
| C2_07370W | 0.00032 | 8.23 | PDB1 | 0.00036 | 18.47 | HEM14 | 0.000339 | 2.38 |
| C1_11200W | 0.00032 | 7.43 | C2_05160C | 0.00035 | 14.69 | SSZ1 | 0.000335 | 3.73 |
| C1_12280C | 0.00032 | 13.05 | C5_04630W | 0.00035 | 16.96 | C3_00850C | 0.000333 | 3.31 |
| RPS6A | 0.00031 | 6.8 | MRPL40 | 0.00035 | 21.3 | PDI1 | 0.000329 | 4.42 |
| C5_04910W | 0.00031 | 8.25 | C2_01740C | 0.00035 | 10.62 | EFT2 | 0.000327 | 4.49 |
| RPS26A | 0.00031 | 8.23 | ACS2 | 0.00034 | 11.98 | MPP10 | 0.000326 | 3.56 |
| DPM1 | 0.00031 | 4.7 | C1_12280C | 0.00034 | 11.64 | NOT4 | 0.000322 | 3.01 |
| MGE1 | 0.00031 | 9.11 | RPT2 | 0.00034 | 15.52 | CR_00490W | 0.000317 | 3.33 |
| PRX1 | 0.0003 | 6.54 | ACH1 | 0.00034 | 8.4 | PES1 | 0.000314 | 1.96 |
| CR_07080W | 0.0003 | 10.9 | TRP99 | 0.00032 | 7.07 | C4_03290W | 0.000306 | 2.33 |
| LPD1 | 0.0003 | 14.16 | PFK26 | 0.00032 | 20.92 | C5_02380W | 0.000305 | 5.34 |
| ARO8 | 0.0003 | 12.5 | KAR2 | 0.00031 | 12.95 | AAF1 | 0.000302 | 3 |
| IMG2 | 0.0003 | 4.44 | CR_07510W | 0.00031 | 8.8 | MIR1 | 0.000297 | 3.3 |
| NSA2 | 0.00028 | 6.98 | PRE8 | 0.00031 | 6.37 | PIL1 | 0.000292 | 6.14 |
| SDH2 | 0.00028 | 9.88 | RHR2 | 0.00031 | 10.24 | C2_05300C | 0.000291 | 3.41 |
| NOP15 | 0.00028 | 5.17 | C1_00160C | 0.00029 | 14 | LSP1 | 0.000286 | 3.85 |
| RFA2 | 0.00027 | 7.2 | NOG2 | 0.00029 | 12.2 | C3_02350W | 0.000286 | 1.78 |
| MGM101 | 0.00027 | 6.75 | C3_01720C | 0.00029 | 10.73 | DED1 | 0.000279 | 2.78 |
| HEM1 | 0.00026 | 7.56 | C3_06760W | 0.00028 | 4.36 | ILV2 | 0.000278 | 5.52 |
| HSP60 | 0.00026 | 4.6 | PGK1 | 0.00028 | 11.51 | DRG1 | 0.000278 | 5.54 |
| ZUO1 | 0.00026 | 7.02 | ALI1 | 0.00028 | 10.04 | IDH2 | 0.000272 | 5.41 |
| NSP1 | 0.00026 | 4.94 | C1_11880W | 0.00028 | 7.14 | C4_02260C | 0.00027 | 5.36 |
| RPT5 | 0.00026 | 15.92 | C2_04570W | 0.00027 | 9.37 | C5_03550W | 0.000256 | 2.85 |
| SUB2 | 0.00026 | 8.76 | GCD2 | 0.00027 | 8.6 | FLO8 | 0.000254 | 1.93 |
| RPD31 | 0.00026 | 8.35 | UBC4 | 0.00027 | 18.37 | C4_00420C | 0.00025 | 2.78 |
| C6_00550W | 0.00025 | 10.57 | PRT1 | 0.00026 | 9.88 | C2_04370W | 0.000232 | 3.8 |
| C4_06680C | 0.00025 | 5.36 | C4_06210C | 0.00026 | 12.55 | PGK1 | 0.000232 | 4.88 |
| C6_02350C | 0.00024 | 4.18 | TKL1 | 0.00026 | 6.06 | PDX1 | 0.00023 | 5.39 |
| SEC24 | 0.00024 | 7.42 | RPN1 | 0.00026 | 11.28 | PIN4 | 0.000216 | 1.39 |
| C1_03280W | 0.00024 | 4.87 | C2_02270C | 0.00025 | 9.03 | C1_08180C | 0.000216 | 2.02 |
| CR_04170W | 0.00024 | 11.4 | RNH1 | 0.00025 | 15.45 | UTP9 | 0.000211 | 3.82 |
| YTM1 | 0.00024 | 4.36 | LAT1 | 0.00025 | 9.64 | ROM2 | 0.000209 | 3.41 |
| SNF12 | 0.00024 | 3.37 | NIP1 | 0.00025 | 9.27 | SER33 | 0.000205 | 3.6 |
| C1_00930C | 0.00024 | 10.42 | ARO8 | 0.00024 | 13.85 | SNF12 | 0.000205 | 4.32 |
| C4_05630W | 0.00023 | 3.28 | TPI1 | 0.00024 | 8.87 | ARO8 | 0.000199 | 1.97 |
| CDC28 | 0.00023 | 7.21 | GSY1 | 0.00024 | 19.52 | KGD1 | 0.000195 | 1.21 |
| LSP1 | 0.00023 | 9.05 | ALD5 | 0.00024 | 14.43 | CAP1 | 0.000192 | 3.6 |
| C7_03000C | 0.00023 | 7.16 | C1_05650W | 0.00023 | 8.73 | C3_00130C | 0.000186 | 1.16 |
| C6_02690C | 0.00023 | 7.8 | C7_03830C | 0.00023 | 8.33 | C1_12030W | 0.000185 | 6.48 |
| ACC1 | 0.00023 | 7.97 | C1_00590W | 0.00023 | 14.08 | YWP1 | 0.000182 | 2.99 |
| C4_00420C | 0.00023 | 4.08 | C1_00040W | 0.00023 | 12.94 | ADE4 | 0.000174 | 3.67 |
| MDJ1 | 0.00023 | 3.1 | C4_04820C | 0.00023 | 8.14 | SRV2 | 0.000172 | 2.01 |
| C5_03980W | 0.00022 | 6.79 | C7_03660C | 0.00023 | 9.84 | MLS1 | 0.000171 | 3.01 |
| GCD2 | 0.00022 | 5.48 | C5_03410C | 0.00023 | 5.2 | CR_02610C | 0.000163 | 2.1 |
| ILV2 | 0.00022 | 3.72 | YST1 | 0.00022 | 17.24 | UBR1 | 0.000161 | 3.2 |
| NOP5 | 0.00022 | 7.08 | VMA6 | 0.00022 | 14.37 | KRE30 | 0.00016 | 3.19 |
| AHA1 | 0.00021 | 7.46 | TAF14 | 0.00022 | 13.31 | C1_01590C | 0.000158 | 2.59 |
| C2_01390W | 0.00021 | 6.66 | C1_01580W | 0.00022 | 6.82 | HIS7 | 0.000157 | 3.12 |
| ADH1 | 0.00021 | 5.66 | LYS22 | 0.00022 | 12.23 | PGA63 | 0.000155 | 2.9 |
| MET3 | 0.00021 | 4.73 | C5_00150C | 0.00022 | 13.63 | SSD1 | 0.000148 | 5.21 |
| YWP1 | 0.00021 | 3.47 | IDH1 | 0.00021 | 11.54 | C6_00170C | 0.000144 | 2.24 |
| FBA1 | 0.00021 | 6.25 | CCT7 | 0.00021 | 12.39 | TKL1 | 0.00014 | 4.93 |
| TIF4631 | 0.00021 | 7.92 | PGI1 | 0.00021 | 5.82 | ACS2 | 0.000139 | 1.05 |
| C1_05630C | 0.0002 | 4.71 | NOG1 | 0.00021 | 6.72 | PAN1 | 0.000136 | 3.19 |
| TCP1 | 0.0002 | 7.27 | RPN12 | 0.00021 | 11.91 | TRP5 | 0.000135 | 2.69 |
| GAR1 | 0.0002 | 5.94 | HSP21 | 0.00021 | 5.29 | C1_00930C | 0.000134 | 1.57 |
| RPG1A | 0.0002 | 4.97 | SAR1 | 0.00021 | 10.53 | UTP13 | 0.00013 | 3.8 |
| C7_02960C | 0.0002 | 6.8 | CEF3 | 0.0002 | 7.14 | NAN1 | 0.000126 | 5.18 |
| PDC11 | 0.0002 | 9.22 | C1_01600W | 0.0002 | 10.24 | C1_04290C | 0.000126 | 2.95 |
| RPL9B | 0.00019 | 10.85 | RPL7 | 0.0002 | 10.85 | NIP1 | 0.000122 | 2.71 |
| VMA5 | 0.00019 | 8.09 | MRPL3 | 0.00019 | 11.39 | RPG1A | 0.000122 | 1.36 |
| MRP7 | 0.00019 | 12.31 | THS1 | 0.00019 | 11.05 | SEC24 | 0.000121 | 1.56 |
| GPD1 | 0.00018 | 3.26 | PIL1 | 0.00019 | 6.82 | SEC27 | 0.000107 | 1.5 |
| C5_00260W | 0.00018 | 3.14 | CR_07320C | 0.00019 | 9.39 | C1_07340W | 0.000107 | 4.59 |
| NSA1 | 0.00018 | 6.83 | RNR21 | 0.00019 | 10.17 | C1_00960C | 0.000102 | 2.98 |
| AAF1 | 0.00018 | 3.19 | PDX1 | 0.00019 | 11.51 | GLT1 | 0.000099 | 1.27 |
| PGK1 | 0.00018 | 4.46 | HIS7 | 0.00019 | 6.06 | SLA2 | 0.000098 | 1.6 |
| CR_10490W | 0.00018 | 3.51 | GLN3 | 0.00019 | 20.67 | SRO77 | 0.000092 | 1.83 |
| LYS22 | 0.00018 | 5.42 | NOP1 | 0.00019 | 5.38 | C1_05630C | 0.000092 | 1.72 |
| C5_01700W | 0.00018 | 6.49 | C2_09660W | 0.00018 | 5.5 | C6_03460W | 0.000091 | 3.62 |
| C1_03790C | 0.00017 | 6.49 | C1_14500C | 0.00018 | 6.36 | RPA135 | 0.000084 | 1.28 |
| ABP1 | 0.00017 | 11.83 | UTP18 | 0.00018 | 8.7 | RGA2 | 0.000082 | 1.15 |
| SSC1 | 0.00017 | 4.09 | C7_03840W | 0.00018 | 6.04 | C1_06550W | 0.00008 | 0.8 |
| C2_09660W | 0.00017 | 4 | UTP9 | 0.00018 | 9.89 | NGG1 | 0.00008 | 2.16 |
| C6_00170C | 0.00017 | 7.41 | C1_10470W | 0.00017 | 12.5 | SPT6 | 0.000079 | 3.07 |
| C1_12610W | 0.00017 | 6.28 | CLC1 | 0.00017 | 7.56 | ECM17 | 0.000077 | 1.8 |
| PRO2 | 0.00017 | 3.87 | FAS1 | 0.00017 | 6.53 | MLP1 | 0.000074 | 1.89 |
| DED1 | 0.00017 | 3.08 | SER33 | 0.00017 | 11.27 | FAS1 | 0.000073 | 2.21 |
| NUP49 | 0.00017 | 3.97 | RPD31 | 0.00017 | 9.72 | C7_00570W | 0.000073 | 1.46 |
| TKL1 | 0.00016 | 5.69 | C6_02880W | 0.00017 | 4.74 | C1_11860W | 0.00007 | 1.56 |
| RPC53 | 0.00016 | 2.96 | URA2 | 0.00017 | 8.71 | C1_03290W | 0.000068 | 3.03 |
| C2_02170W | 0.00016 | 6.37 | FBA1 | 0.00016 | 11.98 | C7_04300W | 0.000061 | 3.03 |
| RPD3 | 0.00015 | 3.6 | CIC1 | 0.00016 | 4.66 | C2_06170C | 0.00006 | 1 |
| PHO23 | 0.00015 | 5.04 | C6_03380W | 0.00016 | 4.35 | C4_01060W | 0.000047 | 0.77 |
| C1_10620W | 0.00015 | 6.7 | C3_00450C | 0.00016 | 8.6 | C5_00190C | 0.000042 | 0.76 |
| C6_02370C | 0.00015 | 10.48 | ZCF29 | 0.00016 | 8.42 |  |  |  |
| C1_09040C | 0.00015 | 5.16 | PMM1 | 0.00016 | 5.56 |  |  |  |
| DIP2 | 0.00015 | 4.78 | UTP5 | 0.00015 | 8.81 |  |  |  |
| VID27 | 0.00015 | 4.78 | SDH12 | 0.00015 | 7.33 |  |  |  |
| CDC19 | 0.00015 | 7.34 | NSA2 | 0.00015 | 5.36 |  |  |  |
| RHR2 | 0.00015 | 4.71 | C2_07220W | 0.00015 | 16.67 |  |  |  |
| C3_05880C | 0.00015 | 5 | NOP15 | 0.00015 | 4.87 |  |  |  |
| FUN12 | 0.00015 | 3.59 | VMA8 | 0.00015 | 7.49 |  |  |  |
| C3_07460W | 0.00014 | 5.95 | C3_04810C | 0.00015 | 7.87 |  |  |  |
| RTG3 | 0.00014 | 3.79 | CR_02420W | 0.00015 | 5.95 |  |  |  |
| C3_00130C | 0.00014 | 2.66 | RPN2 | 0.00014 | 7.14 |  |  |  |
| RVS161 | 0.00014 | 3.32 | CCJ1 | 0.00014 | 7.3 |  |  |  |
| ADE4 | 0.00014 | 3.71 | MRPL10 | 0.00014 | 3.89 |  |  |  |
| C5_04720C | 0.00014 | 3.75 | MYO2 | 0.00014 | 6.73 |  |  |  |
| C3_00850C | 0.00014 | 3.34 | SUB2 | 0.00014 | 11.78 |  |  |  |
| ATP1 | 0.00014 | 2.7 | DPS1-1 | 0.00014 | 8.3 |  |  |  |
| CCT7 | 0.00014 | 2.07 | RAS1 | 0.00014 | 4.14 |  |  |  |
| RPN13 | 0.00013 | 7.19 | PES1 | 0.00013 | 5.79 |  |  |  |
| ALI1 | 0.00013 | 3.67 | CR_00490W | 0.00013 | 4.78 |  |  |  |
| CTA7 | 0.00013 | 2.15 | UTP4 | 0.00013 | 10.09 |  |  |  |
| RAD52 | 0.00013 | 2.38 | PET9 | 0.00013 | 7.97 |  |  |  |
| RET2 | 0.00013 | 14.57 | C3_06700C | 0.00013 | 6.85 |  |  |  |
| C1_13320C | 0.00013 | 3.73 | RPG1A | 0.00013 | 9.66 |  |  |  |
| CR_02610C | 0.00013 | 2.85 | YTM1 | 0.00013 | 7.2 |  |  |  |
| CSI2 | 0.00013 | 4.04 | SEC27 | 0.00013 | 8.84 |  |  |  |
| NIP1 | 0.00013 | 3.27 | C5_00080C | 0.00013 | 10.61 |  |  |  |
| PES1 | 0.00013 | 4.76 | PFK2 | 0.00012 | 5.07 |  |  |  |

|  |  |  |  |  |  |
| --- | --- | --- | --- | --- | --- |
| MIR1 | 0.00012 | 7.95 | CR_10830C | 0.00012 | 11.39 |
| SDS24 | 0.00012 | 7.58 | C2_07290W | 0.00012 | 7.23 |
| RPN1 | 0.00011 | 4.78 | UTP13 | 0.00012 | 6.38 |
| C5_02660C | 0.00011 | 4.29 | CR_03120W | 0.00012 | 4.97 |
| C1_10470W | 0.00011 | 2.38 | HXK2 | 0.00012 | 5.99 |
| FAS1 | 0.00011 | 3.28 | C5_04990W | 0.00012 | 11.04 |
| MSI3 | 0.00011 | 4.33 | C3_07420W | 0.00012 | 5.79 |
| CR_04240C | 0.00011 | 5.02 | CDC48 | 0.00012 | 5.45 |
| TRP5 | 0.00011 | 2.04 | C3_02350W | 0.00012 | 8.55 |
| C3_01850W | 0.00011 | 3.55 | RPA135 | 0.00012 | 4.03 |
| CDC10 | 0.0001 | 4.89 | MSS116 | 0.00012 | 4.64 |
| GZF3 | 0.0001 | 2.44 | RPC40 | 0.00012 | 5.06 |
| SRO77 | 0.0001 | 9.38 | ATP2 | 0.00012 | 6.75 |
| ILS1 | 0.0001 | 9.22 | MRPS9 | 0.00012 | 6.25 |
| IDH2 | 0.0001 | 7.88 | CSH1 | 0.00012 | 5.64 |
| UTP4 | 0.0001 | 1.6 | ILV2 | 0.00011 | 5.56 |
| ACT1 | 9.8E-05 | 5.45 | GRP2 | 0.00011 | 15.84 |
| ARP7 | 9.8E-05 | 5.54 | ARC35 | 0.00011 | 5.49 |
| HOG1 | 9.8E-05 | 2.87 | NMD3 | 0.00011 | 5.19 |
| C5_03000C | 9.7E-05 | 6.45 | C5_00790C | 0.00011 | 4.01 |
| SAM2 | 9.6E-05 | 6.27 | C3_01850W | 0.00011 | 3.42 |
| CR_05550C | 9.6E-05 | 5.36 | CHA1 | 0.00011 | 10.2 |
| C2_03360W | 9.5E-05 | 3.24 | CDC10 | 0.00011 | 5.04 |
| C1_06590C | 9.3E-05 | 11.3 | KRR1 | 0.00011 | 3.63 |
| SLK19 | 9.3E-05 | 2.28 | CHC1 | 0.00011 | 5.18 |
| FLO8 | 9.3E-05 | 1.54 | TCP1 | 0.00011 | 5.42 |
| SEC18 | 9.3E-05 | 4.56 | DRG1 | 0.00011 | 4.08 |
| RDH54 | 9.3E-05 | 2.71 | LYS21 | 0.00011 | 10.05 |
| CDC48 | 0.00009 | 3.64 | AGO1 | 0.00011 | 5.78 |
| C2_04370W | 0.00009 | 2.39 | C6_04100W | 0.00011 | 5.15 |
| CAM1 | 0.00009 | 2.88 | GCN1 | 0.00011 | 4.8 |
| PR26 | 0.00009 | 2.76 | C5_04640C | 0.0001 | 8.52 |
| CR_08290W | 8.9E-05 | 4.61 | ERG6 | 0.0001 | 5.32 |
| SMC1 | 8.9E-05 | 4.04 | ARP7 | 0.0001 | 5.29 |
| RPT4 | 8.6E-05 | 4.5 | HOG1 | 0.0001 | 4.24 |
| DNM1 | 8.5E-05 | 2.12 | HET1 | 9.9E-05 | 10.15 |
| C1_02240W | 8.5E-05 | 4.26 | RNR22 | 9.9E-05 | 4.82 |
| PMA1 | 8.3E-05 | 1.84 | PDA1 | 9.7E-05 | 4.74 |
| CR_03220C | 8.3E-05 | 5.97 | C1_00930C | 9.7E-05 | 4.82 |
| MCM2 | 8.2E-05 | 2.51 | C5_00260W | 9.7E-05 | 3.71 |
| CDC54 | 8.1E-05 | 1.8 | NSA1 | 9.6E-05 | 2.96 |
| SER33 | 0.00008 | 4.55 | LMO1 | 9.5E-05 | 5.54 |
| SEC27 | 7.9E-05 | 2.42 | GLT1 | 9.2E-05 | 4.84 |
| FAS2 | 7.9E-05 | 1.64 | TIF4631 | 0.00009 | 2.12 |
| SEC21 | 7.9E-05 | 4.65 | HSP104 | 8.7E-05 | 3.45 |
| PFK2 | 7.8E-05 | 1.01 | SWD1 | 8.7E-05 | 4.92 |
| SEC26 | 7.8E-05 | 1.94 | SES1 | 8.5E-05 | 6.33 |
| ECM17 | 7.7E-05 | 4.63 | SPT6 | 8.4E-05 | 3.5 |
| C2_04620W | 7.6E-05 | 4.62 | SNF12 | 8.3E-05 | 7.25 |
| PNG2 | 7.5E-05 | 2.67 | C1_12240C | 8.3E-05 | 3.4 |
| PFK1 | 7.5E-05 | 4.8 | MIS11 | 8.2E-05 | 3.91 |
| ATP2 | 7.3E-05 | 0.97 | LPD1 | 7.9E-05 | 10.39 |
| MTS1 | 7.2E-05 | 1.71 | C6_03440W | 7.6E-05 | 4.43 |
| C1_00060W | 7.2E-05 | 2.19 | ARO9 | 7.5E-05 | 6.88 |
| C2_06980W | 7.2E-05 | 2.23 | AIP2 | 7.4E-05 | 3.8 |
| CR_10470C | 0.00007 | 3.36 | HOM3 | 7.2E-05 | 3.13 |
| SLA2 | 0.00007 | 1.22 | PBS2 | 7.2E-05 | 2.94 |
| C4_06210C | 0.00007 | 4.06 | NAN1 | 0.00007 | 3.93 |
| CEF3 | 0.00007 | 6.48 | GAD1 | 6.9E-05 | 4.23 |
| MLS1 | 6.7E-05 | 1.6 | HAS1 | 6.9E-05 | 3.54 |
| UTP18 | 6.7E-05 | 1.59 | CSI2 | 6.7E-05 | 3.42 |
| RPA190 | 6.7E-05 | 2.14 | GGA2 | 6.7E-05 | 2.91 |
| PGI1 | 6.7E-05 | 3.7 | PWP1 | 6.4E-05 | 3.59 |
| PDI1 | 6.6E-05 | 1.16 | C6_02350C | 6.4E-05 | 3.1 |
| C2_02540W | 6.4E-05 | 3.27 | APE2 | 6.3E-05 | 3.79 |
| ELF1 | 6.2E-05 | 3.81 | PGA63 | 6.2E-05 | 4.35 |
| KRE30 | 6.1E-05 | 1.32 | C2_05300C | 6.2E-05 | 5.41 |
| HIS7 | 5.9E-05 | 2.23 | SEC26 | 6.1E-05 | 3.15 |
| C6_04290W | 5.9E-05 | 2.98 | ACC1 | 0.00006 | 3.43 |
| C3_05360C | 5.9E-05 | 8.31 | C1_00640C | 0.00006 | 2.77 |
| TOM1 | 5.6E-05 | 2.34 | C2_06200C | 0.00006 | 1.83 |
| C3_02350W | 5.5E-05 | 3.31 | GLC7 | 5.9E-05 | 4.24 |
| UTP8 | 5.3E-05 | 7.23 | ROB1 | 5.9E-05 | 2.12 |
| GLT1 | 5.2E-05 | 3.42 | DED1 | 5.8E-05 | 5.06 |
| C4_07060W | 5.1E-05 | 2.1 | RGD1 | 5.8E-05 | 2.95 |
| CDC53 | 4.9E-05 | 3.2 | VPS1 | 5.6E-05 | 2.45 |
| CR_03200C | 4.7E-05 | 2.92 | PPH21 | 5.4E-05 | 5.28 |
| NAN1 | 4.4E-05 | 2.72 | PIM1 | 5.4E-05 | 3.25 |
| ERB1 | 4.4E-05 | 3.27 | C1_08110W | 5.4E-05 | 4.95 |
| UBR1 | 4.1E-05 | 2.18 | C1_05630C | 5.3E-05 | 3.25 |
| RPN2 | 3.9E-05 | 2.33 | MSN4 | 5.2E-05 | 2.25 |
| DCK1 | 3.9E-05 | 2.61 | C7_03160W | 5.2E-05 | 2.15 |
| C1_07340W | 3.6E-05 | 2.09 | VMA5 | 0.00005 | 3.87 |
| PIM1 | 3.4E-05 | 2.63 | CR_03200C | 0.00005 | 1.8 |
| C6_03460W | 3.2E-05 | 2.87 | RRP6 | 0.00005 | 5.73 |
| RGA2 | 3.1E-05 | 1.77 | NUP82 | 4.9E-05 | 5.26 |
| C1_11860W | 0.00003 | 1.11 | C2_04370W | 4.8E-05 | 3.41 |
| C1_01590C | 0.00003 | 7.04 | SRP40 | 4.6E-05 | 6.78 |
| PGA63 | 2.9E-05 | 1.49 | C1_03790C | 4.6E-05 | 4.93 |
| SSD1 | 2.9E-05 | 1.8 | SEC12 | 4.6E-05 | 2.26 |
| C1_08180C | 2.8E-05 | 2.44 | DNM1 | 4.5E-05 | 1.73 |
| SPT6 | 2.6E-05 | 1.27 | C1_02240W | 4.5E-05 | 2.17 |
| MLP1 | 0.00002 | 19.05 | PRO2 | 4.4E-05 | 3.59 |
| GCN1 | 1.5E-05 | 0.9 | PMA1 | 4.4E-05 | 5.25 |
| MDN1 | 1.5E-05 | 2.39 | C1_14080W | 4.3E-05 | 3.14 |
| TRA1 | 0.00001 | 1.93 | SHM2 | 4.1E-05 | 2.98 |
|  |  |  | RPD3 | 4.1E-05 | 3.75 |
|  |  |  | C6_02370C | 0.00004 | 2.9 |
|  |  |  | C1_09040C | 0.00004 | 3.29 |
|  |  |  | PNG2 | 0.00004 | 1.83 |

|  |  |  |
| --- | --- | --- |
| CAP1 | 3.9E-05 | 2.81 |
| C1_00060W | 3.8E-05 | 2.73 |
| SRV2 | 3.6E-05 | 2.39 |
| CRM1 | 3.6E-05 | 2.97 |
| PEX1 | 3.6E-05 | 1.74 |
| RPA190 | 3.5E-05 | 2.58 |
| CCT6 | 3.5E-05 | 3.76 |
| C5_02380W | 3.3E-05 | 2.68 |
| PYC2 | 3.3E-05 | 2.63 |
| KRE30 | 3.2E-05 | 1.97 |
| C1_01590C | 3.2E-05 | 3.16 |
| C7_01030C | 3.2E-05 | 1.73 |
| MSI3 | 2.8E-05 | 3 |
| UTP8 | 2.8E-05 | 2.73 |
| C2_00360C | 2.8E-05 | 1.86 |
| CDC53 | 2.6E-05 | 1.86 |
| C4_00420C | 2.4E-05 | 2.47 |
| TOM1 | 2.4E-05 | 1.58 |
| ERB1 | 2.3E-05 | 2.24 |
| SNF2 | 2.3E-05 | 1.18 |
| NAB3 | 2.3E-05 | 2.39 |
| FAS2 | 2.1E-05 | 0.8 |
| SPO72 | 2.1E-05 | 0.7 |
| PFK1 | 0.00002 | 1.82 |
| CDC39 | 1.9E-05 | 0.69 |
| MET10 | 1.8E-05 | 1.46 |
| SSN6 | 1.8E-05 | 1.48 |
| MDN1 | 1.6E-05 | 1.33 |
| C2_02540W | 1.1E-05 | 0.93 |

| Common elements in TAF12L-FLAG TBP-TAP TAF11-TAP : | Common elements in TBP-TAP TAF11-TAP : | Common elements in TAF12L-FLAG TBP-TAP : | Common elements in TAF12L-FLAG TAF11-TAP | TAF12L only | TBP only | TAF11 only |
| --- | --- | --- | --- | --- | --- | --- |
| C1_10620W | TBP1 | RPL43A | NGG1 | NOP10 | C7_01400C | C2_03160C |
| GAR1 | C7_00340C | RPL25 | RPS19A | BBC1 | C4_04600C | RPC10 |
| RPL4B | RIM1 | RPL39 | RPS25B | ACS1 | TRI1 | CMD1 |
| RPL10A | CR_03460W | SSC1 | C4_04390W | ADA2 | CR_04310C | C7_02660C |
| RPL2 | TAF145 | RPL23A | YML6 | CR_10450C | TOA2 | TIF34 |
| RPP2A | TAF4 | RPL32 | C4_02260C | CR_04870C | BRF1 | CR_06800C |
| RPP0 | C2_02500W | RPS15 | C1_05270C | SPT7 | C4_05820W | MNT1 |
| RPL8B | TAF12 | ACT1 | TPI1 | GCN5 | CR_05150W | C4_04330C |
| RPL3 | C3_03930W | RPL28 | C5_00150C | NHP2 | MRPL19 | C1_01860W |
| CR_04450C | SIS1 | TRA1 | NOG1 | SPT20 | PTC2 | C7_00070C |
| RPP1A | C2_10680W | RPS18 | UTP9 | C2_05830C | C4_03090W | C5_02210W |
| RPL10 | HTA1 | RPL13 | C3_00450C | C7_00450C | RFA1 | SGT2 |
| RPL12 | TAF19 | CR_04240C | VMA8 | RPS3 | MCI4 | MP65 |
| SSB1 | TSM1 | RPL35 | CR_00490W | C5_02900W | DCW1 | LAB5 |
| RPS12 | SKP1 | RPL21A | UTP13 | C3_03100C | C1_09620C | C6_00290W |
| RPL15A | C4_04160W | RPS24 | RPA135 | TAF12L | C5_00560W | C3_03410C |
| RPS6A | KGD2 | RPL17B | DRG1 | RPS23A | C1_01680C | CCR4 |
| RPL18 | DBP5 | PR26 | C2_05300C | C6_02310W | C1_01370C | HEM14 |
| RPS8A | C7_00790W | C5_01700W | CAP1 | C3_07050W | C4_06730C | SSZ1 |
| TAF60 | KGD1 | RPL11 | SRV2 | C3_05790C | C7_01210C | MPP10 |
| RPL14 | C1_11200W | HTA2 | C2_00360C | CR_04110W | C4_05900C | NOT4 |
| RPL24A | MGE1 | C2_07680W |  | RPS21B | MAK16 | C4_03290W |
| SBP1 | SDH2 | RPS20 |  | C1_11080W | GSP1 | C5_03550W |
| C2_05710C | HEM1 | C2_05410W |  | RPS10 | CR_10740W_A | PIN4 |
| TAF10 | NSP1 | GFA1 |  | RPS16A | C2_07370W | ROM2 |
| C5_03830C | C4_06680C | IMG2 |  | TEF2 | DPM1 | C1_12030W |
| RPS14B | SEC24 | C1_02330C |  | RPS28B | PRX1 | PAN1 |
| TEF1 | LSP1 | C5_01540W |  | CR_04390C | RFA2 | C1_04290C |
| HHF22 | C6_02690C | RPL20B |  | RPS5 | MGM101 | C1_00960C |
| C1_00180W | MDJ1 | RPT5 |  | C4_03040W | ZUO1 | C1_06550W |
| RPL6 | MET3 | RPL5 |  | C1_03620C | C6_00550W | C7_00570W |
| ERG13 | YWP1 | ATP1 |  | C5_00820W | C1_03280W | C1_03290W |
| C1_00710C | AAF1 | RVS161 |  | MRPL37 | CR_04170W | C7_04300W |
| C2_03560C | C6_00170C | CR_10350C_B |  | RPT6 | CDC28 | C2_06170C |
| RPL19A | NUP49 | C5_01050C |  | CR_10820W_A | C5_03980W | C4_01060W |
| RPL9B | C2_02170W | RPT4 |  | AHP1 | AHA1 | C5_00190C |
| C3_04380C | C3_05880C | C5_04910W |  | C6_02470W | C2_01390W |  |
| RPL27A | RTG3 | C5_02660C |  | UBP8 | C7_02960C_A |  |
| RPL30 | C3_00130C | C2_03950W |  | CR_07220C | GPD1 |  |
| RVB1 | C5_04720C | RPN13 |  | TSA1B | CR_10490W_A |  |
| UBI3 | C3_00850C | SAM2 |  | C6_03430C | ABP1 |  |
| NOP5 | CR_02610C | MRPL40 |  | C1_00110W | RPC53 |  |
| C2_07190C | MIR1 | C2_01740C |  | RPS17B | PHO23 |  |
| MRT4 | TRP5 | RHR2 |  | C3_06970W | DIP2 |  |
| RPS1 | SRO77 | C3_01720C |  | RSM22 | VID27 |  |
| RPP2B | IDH2 | ALI1 |  | RPT1 | FUN12 |  |
| SIK1 | C2_03360W | GCD2 |  | SRB1 | C3_07460W |  |
| HSP70 | C1_06590C | C4_06210C |  | C2_05160C | CTA7 |  |
| RPP1B | SLK19 | RPN1 |  | C5_04630W | RAD52 |  |
| TUB1 | FLO8 | C1_00590W |  | RPT2 | RET2 |  |
| VMA2 | CAM1 | CCT7 |  | ACH1 | C1_13320C |  |
| ASC1 | CR_08290W | PGI1 |  | TRP99 | SDS24 |  |
| TDH3 | ECM17 | CEF3 |  | PFK26 | GZF3 |  |
| MRP7 | MTS1 | RPL7 |  | CR_07510W | ILS1 |  |
| C1_04180W | SLA2 | CR_07320C |  | PRE8 | C5_03000C |  |
| ADH1 | MLS1 | NOP1 |  | NOG2 | CR_05550C |  |
| C1_00900W | PDI1 | C2_09660W |  | C3_06760W | SEC18 |  |
| TIF | C6_04290W | C1_14500C |  | C1_11880W | RDH54 |  |
| C1_05720W | UBR1 | UTP18 |  | UBC4 | SMC1 |  |
| BMH1 | C1_07340W | C1_10470W |  | C2_02270C | CR_03220C |  |
| RPS9B | C6_03460W | RPD31 |  | RNH1 | MCM2 |  |
| EFT2 | RGA2 | FBA1 |  | GSY1 | CDC54 |  |
| RPS21 | C1_11860W | CIC1 |  | ALD5 | SEC21 |  |
| RVB2 | SSD1 | C6_03380W |  | C1_05650W | C2_04620W |  |
| HSP60 | C1_08180C | NSA2 |  | C7_03830C | C2_06980W |  |
| C7_03000C | MLP1 | NOP15 |  | C1_00040W | CR_10470C_B |  |
| GPM1 |  | RPN2 |  | C7_03660C | ELF1 |  |
| RPS7A |  | UTP4 |  | C5_03410C | C3_05360C |  |
| C1_03370W |  | YTM1 |  | YST1 | C4_07060W |  |
| PDC11 |  | CR_10830C_A |  | VMA6 | DCK1 |  |
| C2_04120C |  | CR_03120W |  | C1_01580W |  |  |
| C4_03410W |  | C5_04990W |  | IDH1 |  |  |
| HSP90 |  | RPC40 |  | RPN12 |  |  |
| SSA2 |  | C3_01850W |  | SAR1 |  |  |
| CR_07080W |  | CDC10 |  | C1_01600W |  |  |
| C1_12610W |  | TCP1 |  | THS1 |  |  |
| RPS42 |  | GCN1 |  | RNR21 |  |  |
| C5_00030W |  | ARP7 |  | GLN3 |  |  |
| CDC19 |  | HOG1 |  | C7_03840W |  |  |
| RNR1 |  | HET1 |  | CLC1 |  |  |
| RPS26A |  | C5_00260W |  | C6_02880W |  |  |
| ENO1 |  | NSA1 |  | ZCF29 |  |  |
| CR_00460C |  | CSI2 |  | PMM1 |  |  |
| YDJ1 |  | PWP1 |  | UTP5 |  |  |
| ADE4 |  | C6_02350C |  | C2_07220W |  |  |
| RPS13 |  | SEC26 |  | C3_04810C |  |  |
| SMT3 |  | GLC7 |  | CR_02420W |  |  |
| C4_05630W |  | PIM1 |  | CCJ1 |  |  |
| PDB1 |  | VMA5 |  | MRPL10 |  |  |
| ACS2 |  | CR_03200C |  | MYO2 |  |  |
| C1_12280C |  | SRP40 |  | DPS1-1 |  |  |
| KAR2 |  | C1_03790C |  | RAS1 |  |  |
| C1_00160C |  | DNM1 |  | C3_06700C |  |  |
| PGK1 |  | C1_02240W |  | C5_00080C |  |  |
| C2_04570W |  | PRO2 |  | C2_07290W |  |  |
| PRT1 |  | PMA1 |  | HXK2 |  |  |
| TKL1 |  | RPD3 |  | C3_07420W |  |  |
| LAT1 |  | C6_02370C |  | MSS116 |  |  |
| NIP1 |  | C1_09040C |  | MRPS9 |  |  |
| ARO8 |  | PNG2 |  | CSH1 |  |  |
| C4_04820C |  | C1_00060W |  | GRP2 |  |  |
| TAF14 |  | RPA190 |  | ARC35 |  |  |
| LYS22 |  | MSI3 |  | NMD3 |  |  |
| HSP21 |  | UTP8 |  | C5_00790C |  |  |
| MRPL3 |  | TOM1 |  | CHA1 |  |  |
| PIL1 |  | ERB1 |  | KRR1 |  |  |
| PDX1 |  | FAS2 |  | LYS21 |  |  |
| HIS7 |  | PFK1 |  | AGO1 |  |  |
| FAS1 |  | MET10 |  | C6_04100W |  |  |
| SER33 |  | MDN1 |  | C5_04640C |  |  |
| URA2 |  | C2_02540W |  | ERG6 |  |  |
| SDH12 |  |  |  | RNR22 |  |  |
| SUB2 |  |  |  | LMO1 |  |  |
| PES1 |  |  |  | SWD1 |  |  |
| PET9 |  |  |  | SES1 |  |  |
| RPG1A |  |  |  | C1_12240C |  |  |
| SEC27 |  |  |  | MIS11 |  |  |
| PFK2 |  |  |  | C6_03440W |  |  |
| CDC48 |  |  |  | ARO9 |  |  |
| C3_02350W |  |  |  | AIP2 |  |  |
| ATP2 |  |  |  | HOM3 |  |  |
| ILV2 |  |  |  | PBS2 |  |  |
| CHC1 |  |  |  | GAD1 |  |  |
| PDA1 |  |  |  | HAS1 |  |  |
| C1_00930C |  |  |  | GGA2 |  |  |
| GLT1 |  |  |  | APE2 |  |  |
| TIF4631 |  |  |  | C1_00640C |  |  |
| HSP104 |  |  |  | C2_06200C |  |  |
| SPT6 |  |  |  | ROB1 |  |  |
| SNF12 |  |  |  | RGD1 |  |  |
| LPD1 |  |  |  | VPS1 |  |  |
| NAN1 |  |  |  | C1_08110W |  |  |
| PGA63 |  |  |  | MSN4 |  |  |
| ACC1 |  |  |  | C7_03160W |  |  |
| DED1 |  |  |  | RRP6 |  |  |
| PPH21 |  |  |  | NUP82 |  |  |
| C1_05630C |  |  |  | SEC12 |  |  |
| C2_04370W |  |  |  | C1_14080W |  |  |
| SHM2 |  |  |  | CRM1 |  |  |
| C5_02380W |  |  |  | PEX1 |  |  |
| KRE30 |  |  |  | CCT6 |  |  |
| C1_01590C |  |  |  | PYC2 |  |  |
| CDC53 |  |  |  | C7_01030C |  |  |
| C4_00420C |  |  |  | SNF2 |  |  |
|  |  |  |  | NAB3 |  |  |
|  |  |  |  | SPO72 |  |  |
|  |  |  |  | CDC39 |  |  |
|  |  |  |  | SSN6 |  |  |

**Figure S1.**
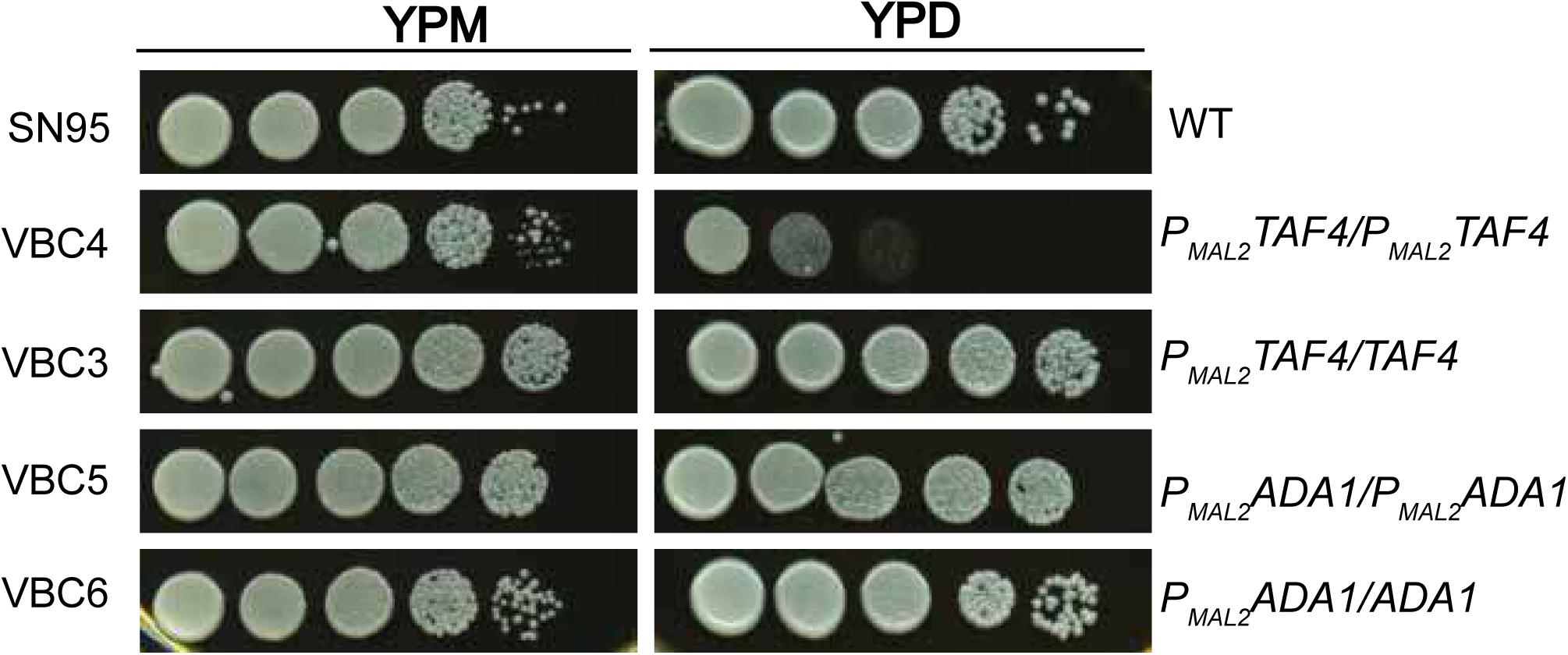
*A*, Growth phenotype analysis of *TAF4* and *ADA1* depleted strains. *A,* Growth phenotype of *TAF4* and *ADA1* depleted strains. Strains were grown in YPM till saturation and serially diluted, spotted on YPM and YPD plates, and incubated at 30°C. Plates were imaged at 36 h.

**Figure S2:**
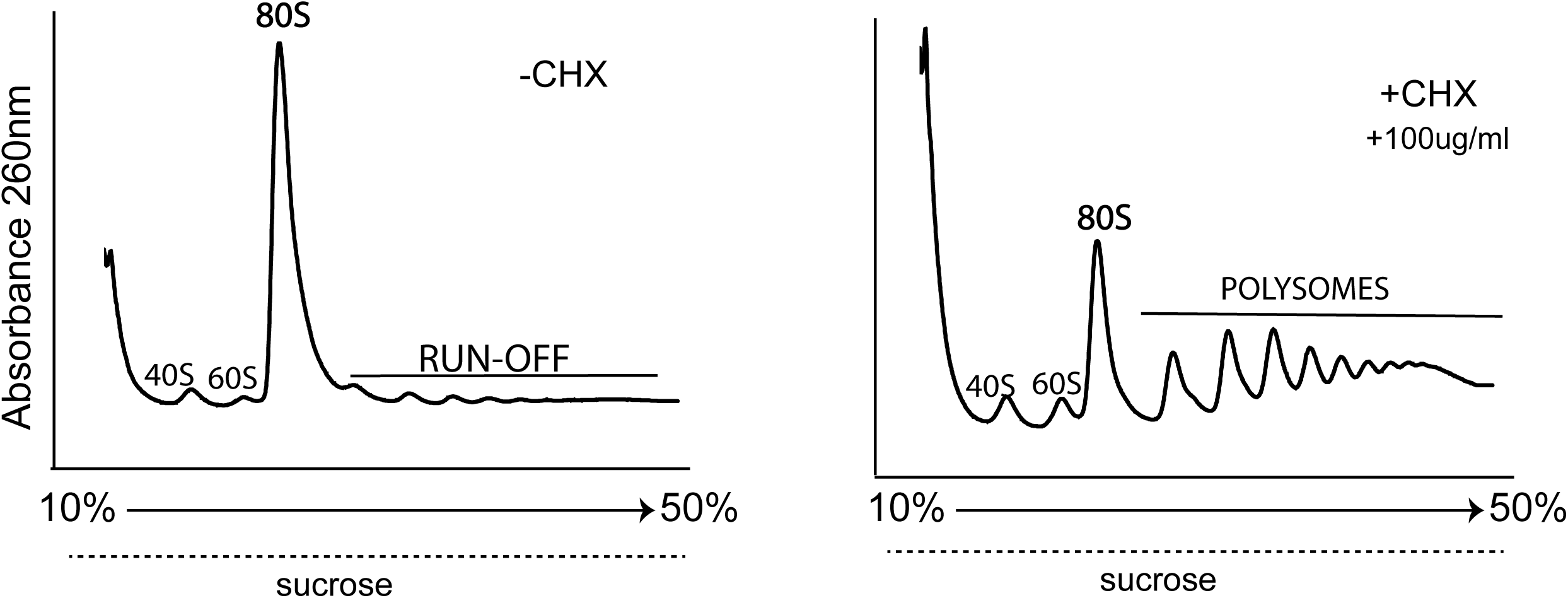
Polysome profiling of *S. cerevisiae*: Cell extracts from untreated(left) or CHX-treated (100µg/ml, right) *S. cerevisiae* BY4741 cultures fractionated through a 10% to 50% sucrose gradient.

## Literature cited

1. Shieh, Y.-W., Minguez, P., Bork, P., Auburger, J. J., Guilbride, D. L., Kramer, G., and Bukau, B. (2015) Operon structure and cotranslational subunit association direct protein assembly in bacteria. Science. 350, 678–680

2. Panasenko, O. O., Somasekharan, S. P., Villanyi, Z., Zagatti, M., Bezrukov, F., Rashpa, R., Cornut, J., Iqbal, J., Longis, M., Carl, S. H., Peña, C., Panse, V. G., and Collart, M. A. (2019) Co-translational assembly of proteasome subunits in NOT1-containing assemblysomes. Nat. Struct. Mol. Biol. 26, 110–120

3. Kamenova, I., Mukherjee, P., Conic, S., Mueller, F., El-Saafin, F., Bardot, P., Garnier, J.-M., Dembele, D., Capponi, S., Timmers, H. T. M., Vincent, S. D., and Tora, L. (2019) Co-translational assembly of mammalian nuclear multisubunit complexes. Nat. Commun. 10, 1740

4. Halbach, A., Zhang, H., Wengi, A., Jablonska, Z., Gruber, I. M. L., Halbeisen, R. E., Dehé, P.-M., Kemmeren, P., Holstege, F., Géli, V., Gerber, A. P., and Dichtl, B. (2009) Cotranslational assembly of the yeast SET1C histone methyltransferase complex. EMBO J. 28, 2959–2970

5. Seidel, M., Becker, A., Pereira, F., Landry, J. J. M., De Azevedo, N. T. D., Fusco, C. M., Kaindl, E., Romanov, N., Baumbach, J., Langer, J. D., Schuman, E. M., Patil, K. R., Hummer, G., Benes, V., and Beck, M. (2022) Co-translational assembly orchestrates competing biogenesis pathways. Nat. Commun. 13, 1224

6. Bernardini, A., Mukherjee, P., Scheer, E., Kamenova, I., Antonova, S., Mendoza Sanchez, P. K., Yayli, G., Morlet, B., Timmers, H. T. M., and Tora, L. (2023) Hierarchical TAF1-dependent co-translational assembly of the basal transcription factor TFIID. Nat. Struct. Mol. Biol. 30, 1141–1152

7. Sinha, I., Kumar, S., Poonia, P., Sawhney, S., and Natarajan, K. (2017) Functional specialization of two paralogous TAF12 variants by their selective association with SAGA and TFIID transcriptional regulatory complexes. J. Biol. Chem. 292, 15587

8. Skrzypek, M. S., Binkley, J., Binkley, G., Miyasato, S. R., Simison, M., and Sherlock, G. (2017) The Candida Genome Database (CGD): incorporation of Assembly 22, systematic identifiers and visualization of high throughput sequencing data. Nucleic Acids Res. 45, D592–D596

9. Grant, P. A., Schieltz, D., Pray-Grant, M. G., Steger, D. J., Reese, J. C., Yates, J. R., and Workman, J. L. (1998) A Subset of TAFIIs Are Integral Components of the SAGA Complex Required for Nucleosome Acetylation and Transcriptional Stimulation. Cell. 94, 45–53

10. Wu, P.-Y. J., Ruhlmann, C., Winston, F., and Schultz, P. (2004) Molecular architecture of the S. cerevisiae SAGA complex. Mol. Cell. 15, 199–208

11. Sinha, I., Poonia, P., Sawhney, S., and Natarajan, K. (2017) Functional specialization of two paralogous TAF12 variants by their selective association with SAGA and TFIID transcriptional regulatory complexes. J. Biol. Chem. 292, 6047–6055

12. Gangloff, Y.-G., Werten, S., Romier, C., Carré, L., Poch, O., Moras, D., and Davidson, I. (2000) The Human TFIID Components TAF _II_ 135 and TAF _II_ 20 and the Yeast SAGA Components ADA1 and TAF _II_ 68 Heterodimerize to Form Histone-Like Pairs. Mol. Cell. Biol. 20, 340–351

13. Werten, S., Mitschler, A., Romier, C., Gangloff, Y.-G., Thuault, S., Davidson, I., and Moras, D. (2002) Crystal Structure of a Subcomplex of Human Transcription Factor TFIID Formed by TATA Binding Protein-associated Factors hTAF4 (hTAFII135) and hTAF12 (hTAFII20). J. Biol. Chem. 277, 45502–45509

14. Juszkiewicz, S., and Hegde, R. S. (2018) Quality Control of Orphaned Proteins. Mol. Cell. 71, 443–457

15. Diebold, M.-L., Fribourg, S., Koch, M., Metzger, T., and Romier, C. (2011) Deciphering correct strategies for multiprotein complex assembly by co-expression: Application to complexes as large as the histone octamer. J. Struct. Biol. 175, 178–188

16. Schwarz, A., and Beck, M. (2019) The Benefits of Cotranslational Assembly: A Structural Perspective. Trends Cell Biol. 29, 791–803

17. Bernardini, A., and Tora, L. (2024) Co-translational Assembly Pathways of Nuclear Multiprotein Complexes Involved in the Regulation of Gene Transcription. J. Mol. Biol. 436, 168382

18. Baumann, K. (2025) Proteins that assemble co-translationally lean on their partner for stability. Nat. Rev. Mol. Cell Biol. 26, 85–85

19. Duncan, C. D. S., and Mata, J. (2011) Widespread Cotranslational Formation of Protein Complexes. PLoS Genet. 7, e1002398

20. Stöcklein, W., and Piepersberg, W. (1980) Binding of cycloheximide to ribosomes from wild-type and mutant strains of Saccharomyces cerevisiae. ANTIMICROB AGENTS CHEMOTHER

21. Shiber, A., Döring, K., Friedrich, U., Klann, K., Merker, D., Zedan, M., Tippmann, F., Kramer, G., and Bukau, B. (2018) Cotranslational assembly of protein complexes in eukaryotes revealed by ribosome profiling. Nature. 561, 268–272

22. Egbe, N. E., Paget, C. M., Wang, H., and Ashe, M. P. (2015) Alcohols inhibit translation to regulate morphogenesis in C. albicans. Fungal Genet. Biol. 77, 50–60

23. Kassem, S., Villanyi, Z., and Collart, M. A. (2017) Not5-dependent co-translational assembly of Ada2 and Spt20 is essential for functional integrity of SAGA. Nucleic Acids Res. 45, 1186–1199

24. Yayli, G., Bernardini, A., Mendoza Sanchez, P. K., Scheer, E., Damilot, M., Essabri, K., Morlet, B., Negroni, L., Vincent, S. D., Timmers, H. T. M., and Tora, L. (2023) ATAC and SAGA co-activator complexes utilize co-translational assembly, but their cellular localization properties and functions are distinct. Cell Rep. 42, 113099

25. Yagita, Y., Zavodszky, E., Peak-Chew, S.-Y., and Hegde, R. S. (2023) Mechanism of orphan subunit recognition during assembly quality control. Cell. 186, 3443–3459.e24

26. Taherbhoy, A. M., and Daniels, D. L. (2023) Harnessing UBR5 for targeted protein degradation of key transcriptional regulators. Trends Pharmacol. Sci. 44, 758–761

27. Singh, R. P., Prasad, H. K., Sinha, I., Agarwal, N., and Natarajan, K. (2011) Cap2-HAP Complex Is a Critical Transcriptional Regulator That Has Dual but Contrasting Roles in Regulation of Iron Homeostasis in Candida albicans. J. Biol. Chem. 286, 25154–25170

28. Green, M. R., and Sambrook, J. (2021) Total RNA Extraction from Saccharomyces cerevisiae Using Hot Acid Phenol. Cold Spring Harb. Protoc. 2021, pdb.prot101691

29. Washburn, M. P., Wolters, D., and Yates, J. R. (2001) Large-scale analysis of the yeast proteome by multidimensional protein identification technology. Nat. Biotechnol. 19, 242–247

30. McDonald, W. H., Ohi, R., Miyamoto, D. T., Mitchison, T. J., and Yates, J. R. (2002) Comparison of three directly coupled HPLC MS/MS strategies for identification of proteins from complex mixtures: single-dimension LC-MS/MS, 2-phase MudPIT, and 3-phase MudPIT. Int. J. Mass Spectrom. 219, 245–251

31. Florens, L., and Washburn, M. P. (2006) Proteomic analysis by multidimensional protein identification technology. Methods Mol. Biol. Clifton NJ. 328, 159–175

32. Eng, J. K., McCormack, A. L., and Yates, J. R. (1994) An approach to correlate tandem mass spectral data of peptides with amino acid sequences in a protein database. J. Am. Soc. Mass Spectrom. 5, 976–989

33. Muzzey, D., Schwartz, K., Weissman, J. S., and Sherlock, G. (2013) Assembly of a phased diploid Candida albicansgenome facilitates allele-specific measurements and provides a simple model for repeat and indel structure. Genome Biol. 14, R97

34. Zhang, Y., Wen, Z., Washburn, M. P., and Florens, L. (2010) Refinements to Label Free Proteome Quantitation: How to Deal with Peptides Shared by Multiple Proteins. Anal. Chem. 82, 2272–2281

